# CALANGO: a phylogeny-aware comparative genomics tool for discovering quantitative genotype-phenotype associations across species

**DOI:** 10.1101/2021.08.25.457574

**Authors:** Jorge Augusto Hongo, Giovanni Marques de Castro, Alison Pelri Albuquerque Menezes, Agnello César Rios Picorelli, Thieres Tayroni Martins da Silva, Eddie Luidy Imada, Luigi Marchionni, Luiz-Eduardo Del-Bem, Anderson Vieira Chaves, Gabriel Magno de Freitas Almeida, Felipe Campelo, Francisco Pereira Lobo

## Abstract

The increasing availability of genomic, annotation, evolutionary and phenotypic data for species contrasts with the lack of studies that adequately integrate these heterogeneous data sources to produce biologically meaningful knowledge. Here, we present CALANGO, a phylogeny-aware comparative genomics tool that uncovers functional molecular convergences and homologous regions associated with quantitative genotypes and phenotypes across species, enabling the fast discovery of novel statistically sound, biologically relevant phenotype-genotype associations. We demonstrate the usefulness of CALANGO in two case studies. The first one unveils potential causal links between prophage density and the pathogenicity phenotype in *Escherichia coli*, and confidently demonstrates how CALANGO supports the investigation of basic causal relationships by enabling a level of counterfactual investigation of observed associations in the data. As a second case study, we used our tool to search for homologous regions associated with a complex phenotypic trait in a major group of eukaryotes: the evolution of maximum height in angiosperms. We confidently identify a previously unknown association between maximum plant height and the expansion of the self-incompatibility system, a molecular mechanism that prevents inbreeding and increases genetic diversity. Taller species also have lower rates of molecular evolution due to their longer generation times, a critical concern for their long-term viability. The new mechanism we report could counterbalance this fact, and have far-reaching consequences for fields as diverse as conservation biology and agriculture. CALANGO is provided as a fully operational R package that can be freely installed from CRAN.

## INTRODUCTION

Living species display a wide range of quantitative variation across both their phenotypes and genomic content. Vast amounts of data describing phenotypic variation and high-quality genomes are available as databases and other structured and unstructured data sources (Groth, et al., 2007; Liolios, et al., 2010). The main bottleneck for the extraction of biologically meaningful knowledge from phenotypic and genomic differences across species, therefore, no longer lies in obtaining such data, but instead on analyzing these heterogeneous data types in a biologically meaningful manner and under a comparative and evolutionary genomics framework to gain insights into the putative genomic mechanisms associated with the evolution of complex quantitative phenotypes (Nagy, et al., 2020).

A major challenge when developing data-modeling schema and statistical workflows to aggregate these data sources is to account for controllable sources of biases and errors, such as data dependences arising from common ancestry (Cornwell and Nakagawa, 2017). Furthermore, most comparative genomics strategies rely exclusively on patterns of variation of homologous genomic elements across genomes as the basic unit of comparison. However, this approach fails to capture molecular functional convergences of non-homologous genomic elements fulfilling the same biological function (International Helminth Genomes, 2019; Tong, et al., 2020). From the computational and statistical perspectives, there is a clear need of a tool capable of not only correcting for multiple hypothesis testing (Hung, et al., 2012), but also mitigating frequent biases in genomic data arising from usual bioinformatics procedures such as genome assembly, gene prediction and annotation (Waterhouse, et al., 2018).

In this article we present CALANGO (*Comparative AnaLysis with ANnotation-based Genomic cOmponents*), a first-principles, general comparative genomics tool designed to account for the aforementioned issues while searching for association between quantitative phenotype/genotypes in distinct species or lineages, and the abundance of annotation terms of their genomic components (Figure 1). These annotations may reflect both homologous regions, as in traditional comparative genomics studies, or molecular convergences based on, e.g., Gene Ontology (GO) terms, which can be used to detect molecular functional convergences in the emergence of complex phenotypes such as sociality and parasitism (International Helminth Genomes, 2019; Tong, et al., 2020).

**Figure 1.**
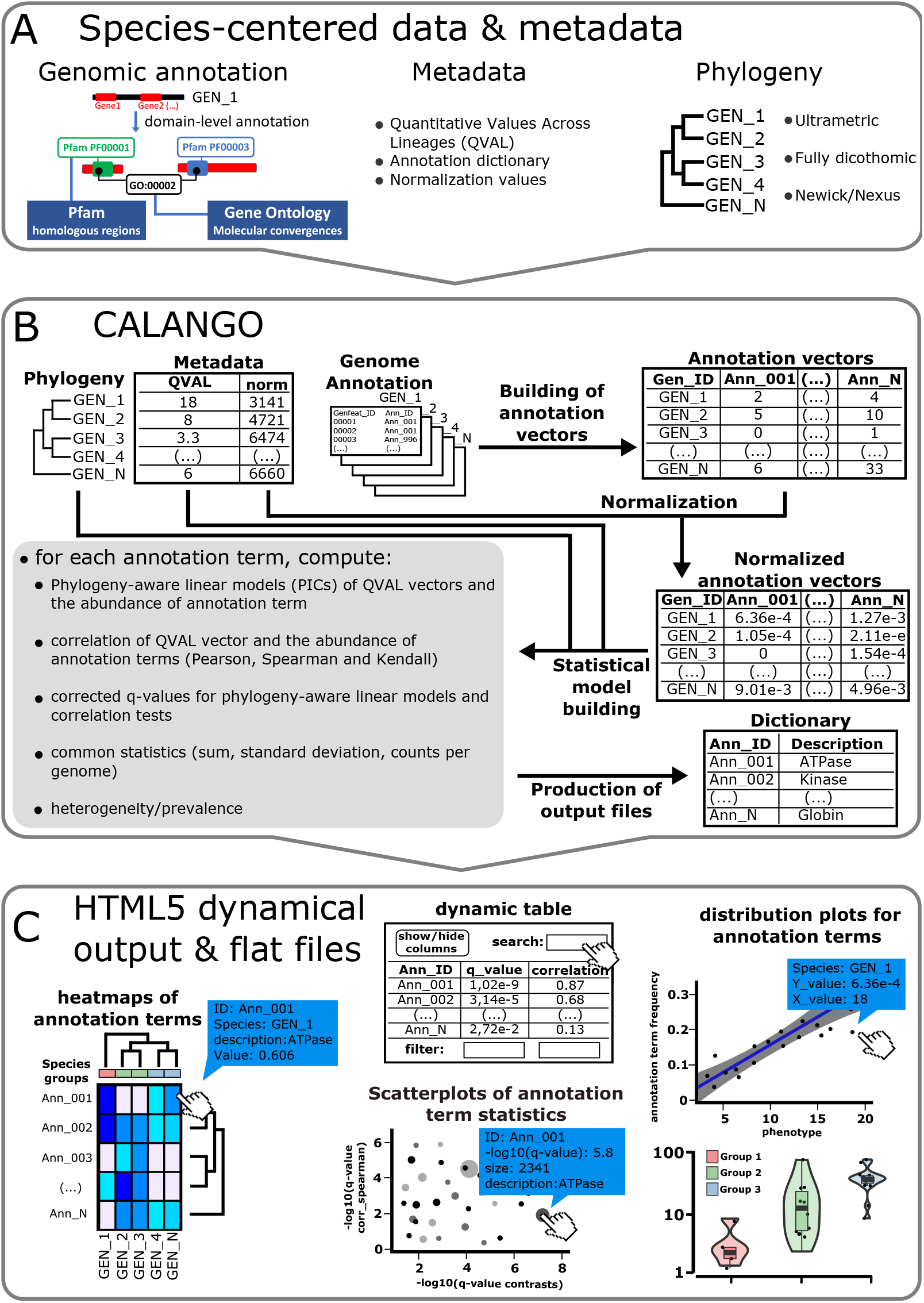
General structure of CALANGO. A) Species-centric input data types needed to run CALANGO: a set of genomic elements from distinct species and their associated annotation data, phylogenetic relationships across species, and the quantitative phenotype/genotype to be surveyed. B) CALANGO execution starts by reading input data and building of annotation vectors (number of occurrences of annotation terms per genome). It can perform optional data normalization to account for relevant variations in genomic content (e.g., genomes with vastly distinct numbers of protein-coding genes). The annotation vectors are then used, together with the quantitative phenotype/genotype, to build phylogeny-aware statistical models and other useful quantities to evaluate possible associations. C) CALANGO outputs dynamical HTML5 reports, which can be accessed using any HTML5-enabled web browser and contain graphical and interactive data representations and summaries, including tables, heatmaps, scatterplots and boxplots highlighting specific aspects of the associations detected.

We validated CALANGO using two complementary case studies that differ in major aspects, such as evolutionary time, taxonomy, and biological phenomena under analysis. The first one comprises the analysis of the biological interaction of integrated bacteriophages (prophages) and *Escherichia coli* lineages, using the density of prophages as a proxy variable. The second evaluated the variation of a complex phenotype in a major group of eukaryotes, namely the evolution of plant height in angiosperms, a key trait for the ecology, physiology and evolution of this group (Mashau, et al., 2021; Zu and Schiestl, 2017).

CALANGO is available as an open-source R package, which can be installed directly from CRAN and also from the project website (https://labpackages.github.io/CALANGO/), where usage examples and long-format documentation can be found. CALANGO outputs interactive HTML5 reports, which facilitate sharing and fast communication of results and integration with existing bioinformatics pipelines.

## MATERIAL AND METHODS

### Genomic data modeling

CALANGO is designed to process distinct classes of genomic components, such as protein-coding genes, protein domains and promoter regions, as well as their respective annotations, to extract biologically meaningful patterns of variation (Supplementary Figure 1A). Our tool explicitly models a genome and the biological functions coded within it by using two associated concepts. The first is a genomic component, defined as a common element type that can be discriminated in the group of genomes under analysis (Supplementary Figure 1A; genomic component IDs in this case could be protein-coding genes, their translated protein sequences, promoters, or protein domains coded within a genome). The second is a set of controlled terms used to annotate the genomic components, which are expected to reflect some specific biological aspect thought to help answer the biological question under investigation, e.g., Pfam domain IDs or GO terms (Supplementary Figure 1A, annotation term IDs). By explicitly dissociating genomic components from their annotation terms, CALANGO makes it possible to use annotation schemas designed to represent the functional similarities of genomic components that may not share a common ancestor, enabling comparative genomics at the function level.

### Input data

Genomic features, annotation, and dictionary files are simple tabular files containing textual information used to describe genomic elements and their annotations (Figure 1A).

- **Annotation/dictionary data:** a Perl script (*calanguize_genomes*.*pl*) that parses Genbank files into high-quality genomic annotation data compatible with CALANGO input data (Figure 1B) is distributed as part of the package. The script performs the following steps:
  A. Downloads genomic data for the species/individual to be analyzed;
  B. Extracts the protein-coding genes described;
  C. Provides a single coding sequence per locus, reporting only the longest coding sequence per locus to avoid possible biases introduced due to the larger number of isoforms described for model organisms (Vogel and Chothia, 2006);
  D. Executes BUSCO for genome completeness evaluation (Waterhouse, et al., 2018);
  E. Annotates all valid protein sequences using InterProScan (Jones, et al., 2014);
  F. Generates CALANGO-compatible files for annotated genomes and dictionaries of annotation terms;
- **Phylogenetic tree data:** CALANGO currently supports fully dichotomous, ultrametric trees in the newick or nexus formats (Figure 1C). Trees with multichotomies are converted into a dichotomous tree with branches of length zero.
- **Metadata:** CALANGO expects a metadata file containing QVAL values, groups for heatmap and boxplot visualization, normalization factors, and other information needed for proper execution (Figure 1A).

### Analyzing data using CALANGO

CALANGO starts its analysis with (i) a set of genomic components from distinct genomes annotated using a common, controlled set of annotation terms; (ii) a dictionary file defining each annotation term in a biologically meaningful way; and (iii) a metadata file containing genome-centered information, such as values for optional normalization of annotation count values in each genome (e.g. total count of annotation terms per genome), and the quantitative phenotype/genotype used to rank genomes (Figure 1A). Users must also provide (iv) an ultrametric phylogenetic tree containing all lineages in a given analysis, which allows CALANGO to correct for phylogeny-related dependencies in the values of annotation terms in distinct genomes and QVAL values (Felsenstein, 1985).

Based on these input files, CALANGO computes different classes of association statistics between the phenotype and annotation terms: three commonly used correlation statistics (Pearson’s, Spearman’s, and Kendall’s correlation values and corresponding p-values) and a phylogeny-aware linear model constructed using Phylogenetic Independent Contrasts (PICs). To account for the multiple hypothesis scenario of simultaneously searching for associations between QVAL and thousands of annotation terms, CALANGO reports FDR-corrected q-values for each association statistic.

CALANGO additionally computes the variance / standard deviation of annotation term counts and two customized statistics that summarize how abundant an annotation term is and how frequently it is observed in distinct genomes: the sum of annotation terms and their prevalence (fraction of genomes where an annotation term is observed).

Two main output structures are provided (Figure 1C). The first is a list-type object containing all computed results, which can be used to survey specific downstream hypotheses. This object also contains all input parameters used to generate the results, therefore providing a simple and convenient way to share results as well as all necessary parameters required for full reproducibility.

The second output is a full interactive HTML5 report / website that can be easily shared, hosted online or browsed locally using any modern web browser. The CALANGO outputs were designed to facilitate more transparent reporting of results and sharing of raw data and code. This user-friendly output facilitates the critical evaluation of all statistics provided by CALANGO in a dynamic tabular and graphical manner.

Four kinds of interactive results are provided by the tool. The first is a biclustered heatmap based on annotation terms (clustered based on their values) and species under analysis (clustered according to the user-provided phylogenetic tree), which allows easy inspection of annotation term distribution across phylogenetic groups and refinement of questions based on interactive exploration of the graph (Figure 1C). The second comprises interactive scatterplots of annotation terms as distributed by their corrected q-values arising from phylogeny-aware models and from common correlation tests. Dot size and transparency are used to highlight interesting annotation terms (both highly frequent and variable across species).

The third type of output is a dynamical table where users may further explore and filter results. Each line contains several computed statistics (e.g., correlation values, q-values for PIC linear model and correlation tests, sum, prevalence, and coefficient of variation) related to a single annotation term, as well as the individual counts of that annotation term in each genome. This table allows users to filter results based on any data column, selecting data slices for further inspection. The dynamic table also contains links to the fourth type of interactive output, namely individual plots of annotation terms results, which includes scatterplots, linear model trend lines and confidence bands for actual data values, ranked data and phylogenetic-aware linear models, and violin plots with superimposed raw data, allowing users to visually inspect how the frequency of annotation terms is distributed in the distinct user-defined groups. The heatmap in Figure 2A and the association scatterplots in Figures 2C and 3C are direct examples of the CALANGO graphical output.

**Figure 2.**
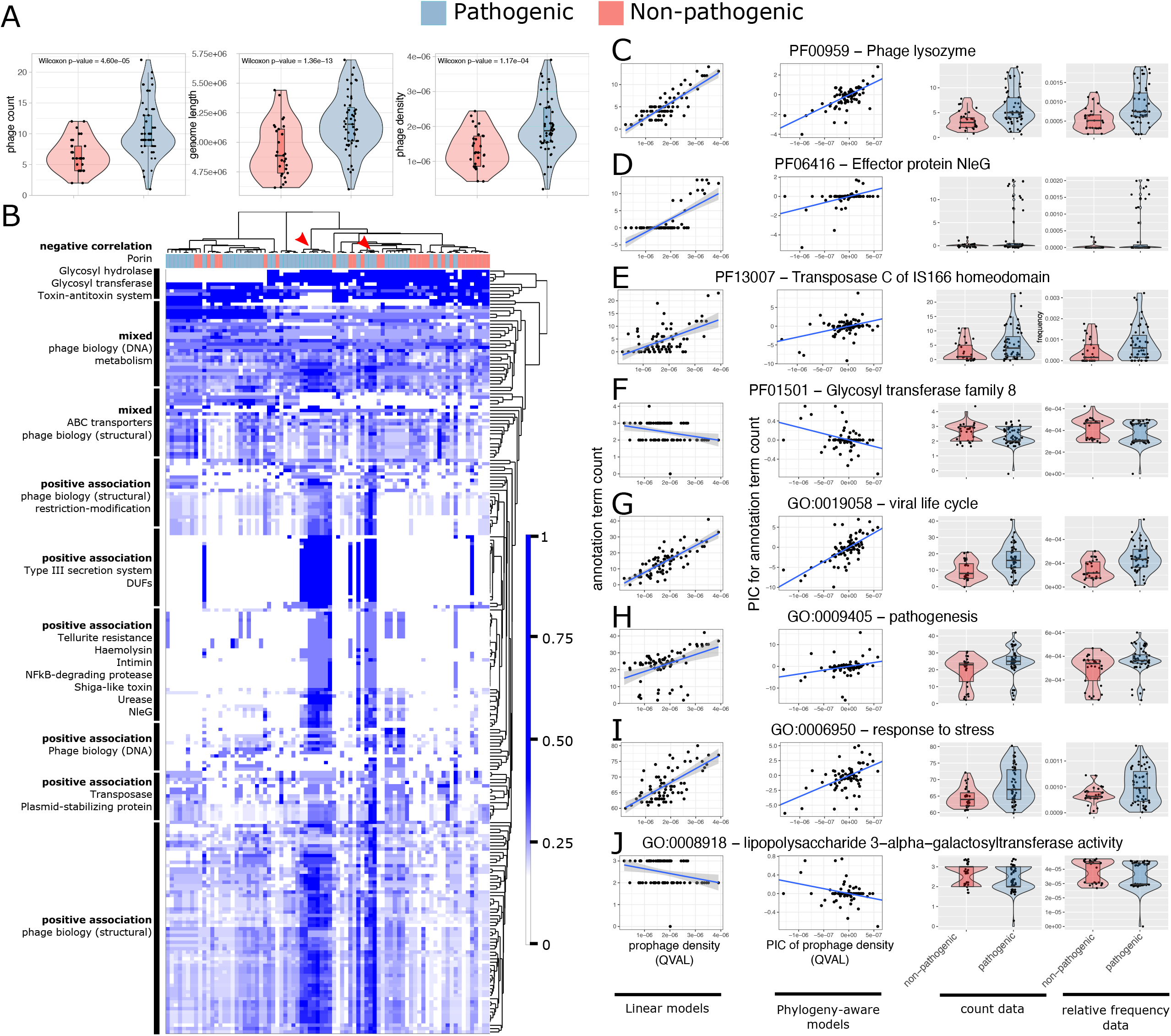
*Escherichia coli* data. A) Comparison of the number of prophages, genome length, and prophage density in pathogenic (blue) and non-pathogenic (pink) *E. coli*. B) Heatmap as produced by CALANGO integrates phylogenetic, phenotypic and annotation data. Species clustering are based on phylogeny, and classes are the same as “A”. Each square contains additional information, and can be accessed through “on mouse over” events. C) Examples of graphical outputs displaying additional information of annotation terms associated with prophage density. From left to right: common association statistics (linear models with Pearson’s correlation data), phylogeny-aware linear models (PIC values for QVAL and annotation data counts) and boxplots with absolute and relative values of annotation terms in the user-defined groups. From top to bottom: examples of annotation terms playing roles in (i) viral life cycle (single domain); (ii) virulence mechanisms (single domain); (iii) transposases (single domain); (iv) LPS biosynthesis (single domain); (v) viral process (GO annotation, multiple domains, functional annotation); (vi) pathogenicity (functional annotation); (vii) stress response mechanisms (functional annotation); (viii) LPS biosynthesis (functional annotation).

**Figure 3.**
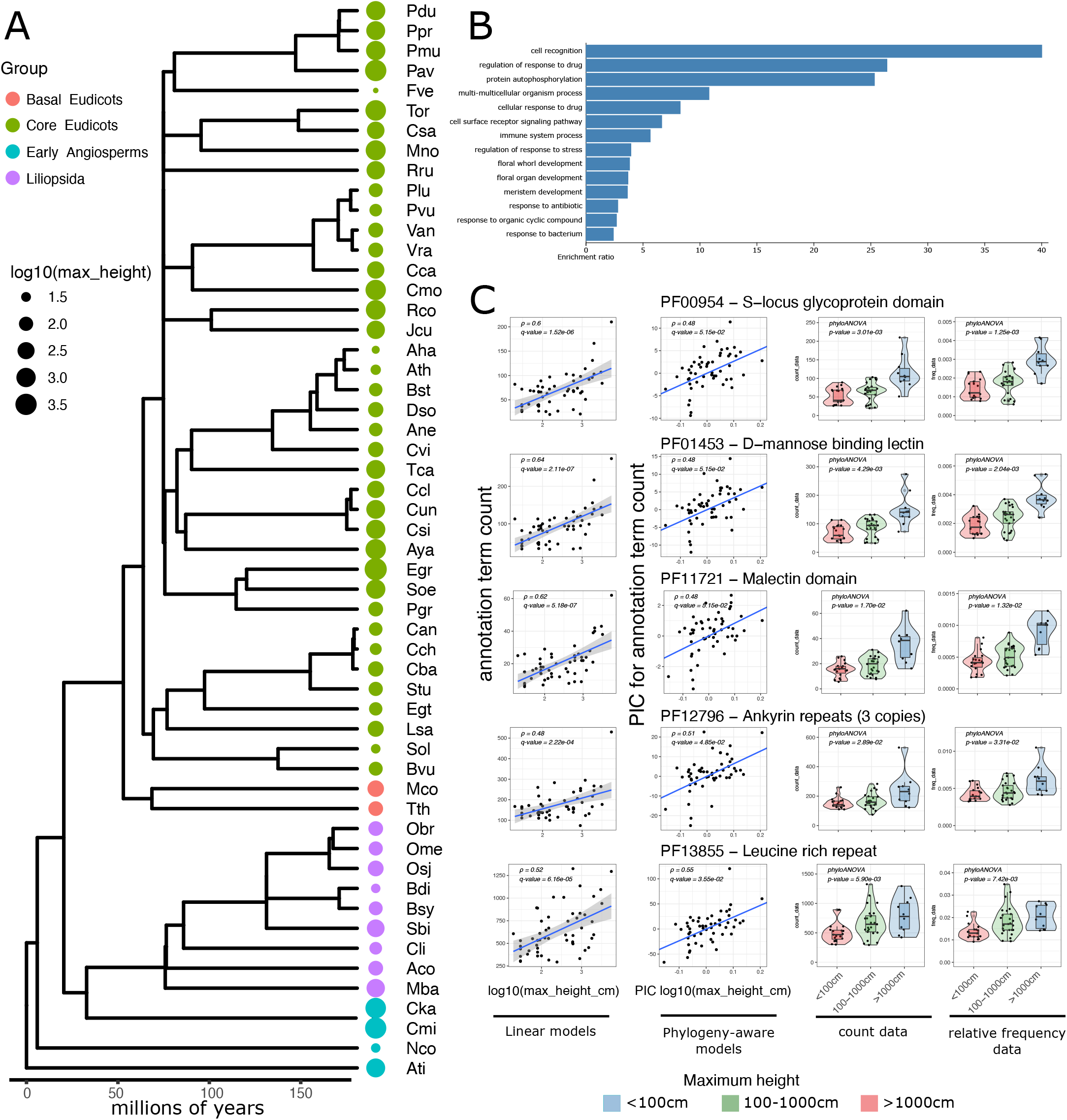
Angiosperm data. A) Maximum height variation across the phylogeny of 54 angiosperms species with high-quality proteomes available. Species names are as follows: *Ananas comosus* (Aco), *Arabidopsis halleri* (Aha), *Arabis nemorensis* (Ane), *Arabidopsis thaliana* (Ath), *Amborella trichopoda* (Ati), *Acer yangbiense* (Aya), *Brachypodium distachyon* (Bdi), *Boechera stricta* (Bst), *Brachypodium sylvaticum* (Bsy), *Beta vulgaris* (Bvu), *Capsicum annuum* (Can), *Capsicum baccatum* (Cba), *Cajanus cajan* (Cca), *Capsicum chinense* (Cch), *Citrus clementina* (Ccl), *Cinnamomum kanehirae* (Cka), *Carex littledalei* (Cli), *Cinnamomum micranthum* (Cmi), *Castanea mollissima* (Cmo), *Cannabis sativa* (Csa), *Citrus sinensis* (Csi), *Citrus* unshiu (Cun), *Cleome violacea* (Cvi), *Descurainia sophioides* (Dso), *Eucalyptus grandis* (Egr), *Erythranthe guttata* (Egt), *Fragaria vesca* (Fve), *Jatropha curcas* (Jcu), *Lactuca saligna* (Lsa), *Musa balbisiana* (Mba), *Macleaya cordata* (Mco), *Morus notabilis* (Mno), *Nymphaea colorata* (Nco), *Oryza brachyantha* (Obr), *Oryza meyeriana* (Ome), *Oryza sativa* (Osj), *Prunus avium* (Pav), *Prunus dulcis* (Pdu), *Punica granatum* (Pgr), *Phaseolus lunatus* (Plu), *Prunus mume* (Pmu), *Prunus persica* (Ppr), *Phaseolus vulgaris* (Pvu), *Ricinus communis* (Rco), *Rhamnella rubrinervis* (Rru), *Sorghum bicolor* (Sbi), *Syzygium oleosum* (Soe), *Spinacia oleracea* (Sol), *Solanum tuberosum* (Stu), *Theobroma cacao* (Tca), *Trema orientale* (Tor), *Thalictrum thalictroides* (Tth), *Vigna angularis* (Van). B) Biological processes significantly enriched in protein-coding genes containing domains associated with maximum height in angiosperms. C) Examples of protein domains significantly expanded in taller plants fulfilling different biological roles. From top to bottom: (i) and (ii) increase of genetic diversity through cross-pollination; (iii) cell wall biology; (iv) embryogenesis; (v) immunity and stress response mechanisms. From left to right: linear models from raw count data, phylogeny-aware linear models; boxplot with raw counts; boxplot with normalized counts.

### Experimental design for case studies

Two annotation schemas (Supplementary Figure 1B) were used to annotate the same genomic elements in each case study, allowing us to contrast the results and evaluate the influence of annotation schema in the analyses:

#### (domain2Pfam)

A domain-level analysis, in which individual predicted protein domains were annotated using a traditional homology-based annotation schema (Pfam IDs) to emulate the results obtained through standard comparative genomics analyses;

#### (domain2GO)

The same domain-level genomic components were annotated using the GO term annotation for the Pfam domains, to highlight how CALANGO may identify functional convergences of non-homologous sequences not detectable in the first analysis.

Finally, as we are performing exploratory data analyses aimed at both validating CALANGO and finding associations that are potentially useful for downstream hypothesis generation, we used a q-value cutoff of < 0.1 for the selection of significative associations.

### Acquiring data for *Escherichia coli*

A thorough literature review was performed to select *E. coli* genomes that could be unambiguously classified as pathogenic or non-pathogenic according to GOLD metadata (Supplementary Table 1) (Mukherjee, et al., 2017). We evaluated genome assembly quality based on the expected gene content using BUSCO 3.0 and the *Enterobacteriales* order (Waterhouse, et al., 2018). All genomes have BUSCO values of single-copy orthologs greater than 0.95. We predicted the numbers and location of prophages in *E. coli* chromosomes using PHASTER with standard parameters, and considered all classes of prophages (Arndt, et al., 2016).

To obtain an ultrametric phylogenetic tree for the 80 *E. coli* genomes we initially computed groups of homologous genes using OrthoMCL v2 (Fischer, et al., 2011) with default parameters and DIAMOND (Buchfink, et al., 2015) for the sequence alignment step. At this point, we included two species from the same family as external groups: *Escherichia fergusonii* (ATCC 35469, NC_011740) and *Shigella dysenteriae* (Sd197, NC_007606) (Chaudhuri and Henderson, 2012). We proceeded by randomly selecting 25 universal 1-1 orthologs from OrthoMCL output file to be used for downstream phylogenetic analyses (Supplementary Table 1).

The global alignment of coding regions was done using MUSCLE (Edgar, 2004), followed by the manual curation of alignments using MEGA X (Kumar, et al., 2018). We used SequenceMatrix (Vaidya G., 2011) to concatenate individual alignments in a single matrix using, and MEGA X to select the best nucleotide substitution models for individual gene alignments. We used MRBAYES (Pang, et al., 2015) as provided by the CIPRES SCIENCE platform to estimate phylogenetic relationships based on the Bayesian information criterion (BIC) and assuming a minimal mutation rate (µ) of 2×10^−10^ mutations per nucleotide site per cell division, following the work of Lee *et al*. (Lee, et al., 2012). The parameters of the posterior probability model, tree branch lengths, and tree topology were obtained using the *Metropolis-coupled Markov Chain Monte Carlo* (MCMCMC) algorithm. Two independent runs of two simultaneous chains for 20 million generations were performed for each analysis, collecting tree samples every 500 generations and discarding the 20% first runs as a burn-in step. Tracer v1.6 was used to evaluate MCMCMC convergence and the conjoint distributions of parameters and trees (Rambaut, et al., 2018). The final summarized tree topology was saved in *newick* format and used as input for our analyses.

### Removal of genes of viral origin in *E. coli* genomes

For each protein-coding gene in all 80 *E. coli* genomes, we generated a BED file containing the IDs of coded proteins and the genomic coordinates of their corresponding genes. We proceeded by generating BED files for each predicted integrated prophage from the output of PHASTER. We then used bedtools (Quinlan, 2014) to remove all protein-coding genes located within prophage coordinates, generating simulated bacterial genomes where genes of viral origin were removed and blocking out any effect of these genes in the CALANGO output while holding all other variables constant, which allowed us to use CALANGO as a support tool to investigate potential causal relationships.

### Manual curation of Pfam domains associated with prophage density

We used the individual description of Pfam IDs provided by InterProScan together with the Pfam ID entries from the Pfam website (El-Gebali, et al., 2019) to classify all Pfam IDs associated with prophage density to the broadest categories that functionally encompass them (Supplementary Table 2, sheet “domain2PfamCount”, column “major_role”).

### Acquiring data for Angiosperms

We downloaded the predicted proteomes of species of flowering plants from NCBI(O’Leary, et al., 2016) and Phytozome(Goodstein, et al., 2012). We detected a high level of duplication in most proteomes due to the presence of protein isoform sequences produced by variant mRNA splicing. Two methods were applied to reduce the redundancy arising from the presence of isoforms. The first was to summarize the NCBI proteomes per genomic locus, keeping only the longest protein sequence for each gene based on the “locus_tag” or “gene_id” information. As proteomes from other databases do not contain these fields in their sequence headers, we used the CD-HIT software with the identity threshold set to 1.0 to reduce the redundancy level for those proteomes (Li and Godzik, 2006).

We used BUSCO to assess the assembly quality of the proteomes based on the expected content of nearly universal ortholog genes (Embryophita db10) (Waterhouse, et al., 2018). Only proteomes with completeness higher than 80% and the rates of duplicated and fragmented genes lower than 15% were kept. We obtained the species phylogenetic relationship from the Time Tree of Life database (Kumar, et al., 2017), and used InterProScan (Jones, et al., 2014) to perform a *de novo* annotation using Pfam data on the predicted proteomes. A total of 54 species of Angiosperms were selected as having the three data types needed to execute CALANGO: high-quality non-redundant proteomes, available quantitative phenotype (maximum plant height) and phylogenetic mapping (Supplementary Table 3).

A considerable fraction of angiosperms is polyploid, and may also have been subjected to previous whole-duplication events (Ren, et al., 2018). These facts may introduce biases when using count data alone, as we have done for the *E. coli* data, since a much more variable genomic content is expected in the non-redundant proteomes of the flowering plants (even though our filtering pipeline removed genomes with high gene duplication values). CALANGO may use a normalizing value (e.g. the total number of domains found in a non-redundant proteome) to compute relative frequencies of annotation terms. However, as we also demonstrated, this metric may introduce spurious associations (See Supplementary File 1, section “Evaluating annotation term frequencies and counts” and Supplementary Figure 2 A-B for a deeper discussion on the biases of using relative frequencies and absolute counts).

To account for all these facts, annotation terms were considered to be associated with the maximum plant height only if both the relative frequency and absolute count were found to be significantly associated. We used CALANGO to search for significant associations with the following cutoffs: 1) q-value for corrected contrasts and Pearson’s correlation < 0.1; and 2) annotation terms with total occurrence count > 50 (to search for major expansions and prevent spurious associations due to the common errors and pitfalls found in eukaryotic genomes).

For the comparative genomics analysis, we cross-referenced genomic annotation data, phylogenetic information and the logarithm of the plants’ maximum height values to search for Pfam domains and GO terms associated with this phenotype using the *domain2Pfam* and *domain2GO* annotation schemas (Supplementary Figure 1B).

### Estimation of ancestor states for heigh in Angiosperms

We used the method of maximum likelihood implemented in the “fastAnc” function of the phytools package to estimate the ancestral height state for all internal nodes of the phylogeny, based on the height values for the 54 extant species under analysis (Revell, 2012).

## RESULTS

### CALANGO - a brief overview

CALANGO provides a flexible platform for investigating the variation of quantitative phenotypes/genotypes in a phylogeny – from now on referred as Quantitative Values Across Lineages (QVALs) – using current genomic knowledge (Figure 1). By dissociating genomic components from their annotation schemas, CALANGO allows comparative genomics analyses at several levels, such as promoters, domains or genes; and, by using distinct annotation schemas, CALANGO enables the search for associations of both homologous regions and functional molecular convergences as provided by, e.g., GO annotation (Supplementary Figure 1A; Methods, section “Genomic data modeling”).

Starting with a set of species/linages and their proper data types, CALANGO searches for associations of annotation terms and QVALs using comparative methods to deal with the data dependencies that emerge when analyzing phylogenetically related organisms, together with traditional statistical models (Figure 1B; Methods, section “Analyzing data using CALANGO”). It also allows normalizing the relative abundance of annotation terms in different genomes by, e.g., genome length or total number of protein-coding genes, to account for the high variability in genome sizes and contents (Supplementary File 1, “Evaluating annotation term frequencies and counts”).

CALANGO outputs a dynamic HTML5 report containing graphical elements and tables with association statistics and other useful quantities, intended to instigate users to interact and actively interpret the results. The full set of user-defined input parameters is also returned alongside all computed results as a *list* object, which can be easily integrated into other bioinformatics pipelines (Figure 1C).

### Case study 1: coevolution of *Escherichia coli* lineages and their integrated bacteriophages

*Escherichia coli* have a remarkable genomic variability, with a considerable fraction of this variation comprising horizontally transferred genes through integrated bacteriophages (prophages) (Touchon, et al., 2009). This genomic diversity is reflected in the distinct ecological niches occupied by this bacterium, which is found in several body niches of animal hosts as a commensal or pathogen. Bacteriophage infections are not always deleterious to their bacterial hosts. While obligate lytic phages represent agents of cell death and population control, persistent lysogenic phages are responsible for gene transfer and mutualism. In a microbial population the lysis-lysogeny events are dynamic and extremes of a continuum comprising antagonistic and beneficial biological interactions (Correa, et al., 2021).

Virulence factors are an archetypal example of bacteriophage-mediated horizontal gene transfer that can result in fitness increase in new bacterial hosts, including pathogenic *E. coli* (Steyert and Kaper, 2012). Despite being a well-known phenomenon, we are not aware of any systematic evaluation of the association between prophage occurrence and the abundance of non-homologous virulence factors. As prophages are themselves genomic elements with specific coordinates, this case study also allows us to selectively remove the effect of viral genes on CALANGO’s results, enabling the potential investigation of associations of causal origin. Therefore, we consider this as an interesting scenario to evaluate our tool, as it has expected causal associations and represents a complex biological interaction, likely to contain previously undescribed biological phenomena.

We performed a thorough literature review to select 80 *E. coli* lineages with both gapless genomes (plus plasmids, when available) and reliable information regarding its pathogenicity status, and computed the annotation, phylogenetic and QVAL data needed to run CALANGO (Supplementary Table 1). Pathogenic *E. coli* were found to have a significantly higher number of prophages, even after accounting for variations in genome size (Figure 2A; Supplementary File 1, section “Integrated bacteriophages in pathogenic and non-pathogenic *E. coli*”). We proceeded by using the prophage density as a proxy QVAL to further survey this biological interaction.

To objectively evaluate GO usefulness to detect molecular functional convergences, we annotated the same set of protein domains found in each bacterial genome by using either their Pfam IDs (*Pfam2domain*) or the GO terms associated with them (*Pfam2GO*) (Supplementary Figure 1B, Methods, section “Experimental design for case studies”). We found GO annotations to be more abundant and more prevalent than our *Pfam2domain* annotation, a scenario compatible with the integration of biological functions from non-homologous domains at the GO space (Supplementary File 1, section “Comparison of functional and homology-based annotation schemas”; Supplementary Figure 1C).

### Homologous regions and biological roles associated with prophage density in *E. coli*

We found 230 out of the 3,335 Pfam domains observed in our *E. coli* dataset (6,8%) to be significantly associated with prophage density (corrected q-values for both phylogeny-aware models and Pearson’s correlation < 0.1). Figure 2B is a heatmap of these terms as produced by CALANGO together with our manual annotation. Of these, 207 domains presented positive correlation (from 0.28 to 0.85) and 23 showed negative correlation (from -0.26 to -0.47) (Supplementary Table 2, sheet “domain2PfamCount”; see also Supplementary File 1, section “Evaluating annotation term frequencies and counts” and Supplementary Figure 2A-B for the rationale of working with raw count data instead of normalized frequencies).

We found the majority of positively associated domains (125/207 domains, 60.4%) to have clear roles in viral life cycle, such as lysozymes and integrases (Figure 2B). Figure 2C illustrates a typical CALANGO output for one of these domains (with additional examples in Supplementary Figure 3A-C). The second largest category encompasses several classes of virulence factors (58/207 domains, 28.0%) (Figure 2B; Figure 2D; Supplementary Figure 3D-E). Some virulence factors such as Shiga-like toxins and effectors of the Type III secretion system are known to be commonly horizontally transferred by bacteriophages in specific *E. coli* pathotypes (Ehrbar and Hardt, 2005; Steyert and Kaper, 2012), an association also detected by CALANGO (Supplementary File 1, section “Homologous genes and biological roles associated with prophage density in *E. coli*”). These two categories provide a compelling example of how CALANGO can uncover known associations of causal origin. Some domains of unknown function (DUFs) have a distribution pattern similar to that of virulence factors (Figure 2B), suggesting these DUFs may be uncharacterized pathogenicity domains and demonstrating how CALANGO can be used to prioritize targets for experimental characterization.

CALANGO also detected positive associations that unveil novel biological interactions between immune genes found in bacterial genomes, prophages, and other classes of mobile elements such as transposases and plasmids. Furthermore, several of these homologous regions are located outside prophage regions, suggesting a complex interplay of symbiosis and competition between them (Figure 2B; Figure 2E; Sup. Figures 3F-H; see also Supplementary File 1 for a deeper discussion on these associations).

The 23 negative associations suggest that *E. coli* lineages with fewer integrated prophages – which are also less likely to be non-pathogenic – have a set of genes enabling a greater diversity of lifestyles at several levels, ranging from metabolic pathways and membrane transport to community-level processes, such as biofilm formation (Figure 2B; Supplementary Figure 3I-L). Interestingly, we observed negative associations of prophage density and components of the cell wall and of the LPS biosynthesis pathway (Figure 2B; Figure 2F). Both classes of molecules are receptors of bacteriophages for cellular infection, but also major activators of the vertebrate immune system (Bertozzi Silva, et al., 2016; Park and Lee, 2013; Wolf and Underhill, 2018). These undocumented negative associations may be a consequence of the selective pressure against prophage infection resulting in the loss of these components. However, these losses can also confer an advantage to pathogenic bacteria when infecting vertebrate hosts, as they may be less likely to trigger host’s immune responses, and could represent a previously unknown aspect of the emergence of a virulence phenotype in this species.

The results provided by the *domain2GO* schema largely support the same conclusions found by our manual curation of the *domain2Pfam* results for both positive and negative associations, highlighting how GO annotation provides an interpretability that is at least qualitatively equivalent to human curation (Figure 2G-H; Supplementary Figure 4A-N; Supplementary Table 2, sheet “domain2GOCount”). Additionally, as several of these biological roles are performed by non-homologous domains (Supplementary File 1, section “GO annotation provides curation-level of biological knowledge”), these results further demonstrate how CALANGO, together with a GO annotation, allows comparative genomics analysis at the function level.

The prophage-mediated horizontal transfer of virulence factors is a known mechanism for fitness increase of pathogenic *E. coli* (Steyert and Kaper, 2012). We also found a positive association of the term GO:0006950 (*response to stress*, Pearson’s correlation of 0.69, q-value for contrasts of 4.65e-05), which may represent a new example of virus-mediated transfer of non-homologous fitness genes that fulfill a common biological role (Figure 2I; Supplementary File 1, section “Stress response genes are associated with prophage density in *E. coli*”; Supplementary Table 2). We hypothesize that integrated prophages could also contribute to fitness increase of bacteria specifically under conditions of stress, such as the host’s immune response against pathogenic bacterial lineages (Wang, et al., 2010), providing yet another dimension of this complex biological interaction and providing additional evidence of how GO-level annotation can support the detection of previously unknown associations.

Since prophages are genomic elements with defined genomic coordinates within bacterial genomes, this case study allows us to perform an *in silico* controlled experiment to evaluate the ability of CALANGO to support the investigation of a potential causal relationship, namely that annotation terms are associated with prophage density *because* they annotate protein-coding genes of viral origin located within prophage coordinates (Supplementary File 1, section “Associations with prophage density after removal of genes of viral origin”).

This experiment consisted of re-executing CALANGO holding the QVAL values fixed (i.e., as calculated previously) and removing all genes of viral origin. This effectively blocks out possible effects of prophage genes on the output of CALANGO, allowing us to test whether the significant associations detected earlier between the annotation terms and the QVAL are indeed due to genes of viral origin.

As expected, the vast majority of protein domains annotated as being of viral origin were no longer significantly associated with prophage density after the removal of genes of viral origin (123/125, 98.4%). A similar pattern was found for GO terms (Supplementary Table 2, sheets “domain2PfamCountLessPhages” and “domain2GOCountLessPhages”). Interestingly, several classes of virulence factor domains were still significantly associated with prophage density after the removal of genes of viral origin, a scenario compatible with bacteriophage-mediated horizontal gene transfer followed by prophage degeneration. However, several other classes virulence domains that were totally or mostly located within detectable prophage genomes were not found to be associated after blocking the effect of viral genes, which suggests a synergistic interaction between virulence factors acquired from different evolutionary origins (Supplementary Table 2, sheet “virulence_factors”).

This *in silico* manipulative experiment showcases the ability of CALANGO to support the investigation of basic causal relationships by enabling a level of counterfactual investigation of observed associations in the data. While this is still short of a fully developed causal inference package for genomic data, the ability to uncover some causal relationships from data by *in* silico isolation and testing of the influence of possible confounders can provide valuable insights into biologically meaningful phenomena, as illustrated in this case study.

### Case study 2: Homologous regions associated with maximum height in Angiosperms

Maximum height is a key trait in the ecology, physiology, and evolution of land plants, and the understanding of the molecular mechanisms associated with the emergence of complex phenotype has consequences for fields as diverse as conservation biology and agriculture (Falster and Westoby, 2003; Moles, et al., 2009). Angiosperms, or the flowering plants, are the largest and most diverse group of land plants, displaying a remarkable phenotypic variation, including in plant height. Artificial selection experiments within single species supports the notion that plant height is a trait strongly controlled by genes that can evolve fast under phenotypic selection, with h^2^ values ranging from 0.5 to 0.9 (Peiffer, et al., 2014; Zu and Schiestl, 2017). However, there is a lack of studies associating the evolution of this phenotype across species under a comparative genomics perspective.

Increases in height seems to be associated with reproductive success, a relationship thought to be caused by several factors, including increased pollination, seed dispersal mechanisms, and access to light. Thus, tall plants that emerged from short ancestors likely experienced positive selection for height, a trait that is potentially under selection in natural populations (Zu and Schiestl, 2017). Shorter plants, on the other hand, have smaller generation times and, consequently, higher rates of evolution, which provides a higher capacity of phenotypical change in distinct environments, and is a concern for the long-term viability of taller species and the ecosystems that depends on them (Lanfear, et al., 2013).

The evolution of a complex phenotype like height is likely coupled with many other plants traits, such as rates of mitosis in meristematic tissues, cell expansion, development of leaves and reproductive organs, pollination syndrome, community composition, among others, and plays important roles in the success of establishment of distinct species (Mashau, et al., 2021; Moles, et al., 2009). As such, the evolution of height in Angiosperms represents a compelling case study to further evaluate CALANGO and demonstrate its usefulness to reveal new biological knowledge.

### Protein domains associated with maximum height in flowering plants unveil independent expansions of reproductive and developmental processes in taller species

We surveyed the specialized literature and sequence databases, together with our *in-house* annotation pipeline, to gather the annotation, phylogenetic and QVAL information for 54 angiosperms species with high-quality non-redundant proteomes available (Supplementary Table 3, see Methods, section “Acquiring genomic, phylogenetic, and phenotypic data for Angiosperms”). Our dataset has species with maximum height varying from 20 cm (the wild strawberry *Fragaria vesca*, Rosaceae) to 55 meters (the tree *Eucalyptus grandis*, Myrtaceae), more than two orders of magnitude (Figure 3A, Supplementary Figure 5A). We found the ancestor states of height in Angiosperms to be highly uniform, with internal nodes having mostly average values and phenotypic extremes occurring multiple times, a pattern compatible with independent emergence of this trait (Figure 3A, Supplementary Figure 5B).

We again used the *domain2Pfam* and *domain2GO* annotation schemas to search for homologous regions and biological roles associated with maximum height. From a total of 4066 domains with at least 50 copies when considering all species, we found seven of them to be positively associated with increase in the plants’ maximum height (Supplementary Table 4, sheet “associated_domains”, see also Methods, section “Experimental design for case studies” for the parameters, experimental design and rationale).

Even though no GO term was found to be significantly associated with the QVAL, an enrichment analysis using the 457 *Arabidopsis thaliana* genes annotated to these Pfam domains found an overrepresentation of genes belonging to reproduction, embryogenesis pathways, including secondary growth, immune system, and stress response mechanisms (Figure 3B-C; Supplementary Table 4, sheet “Arabidopsis_genes”). The increase in the number of genes coding for some of these processes, such as immunity and wood tissue development, has already been reported for *Populus trichocarpa* - a model organism for tree biology - when compared with *A. thaliana* (Tuskan, et al., 2006). The expansions of immune and stress response genes are likely to represent adaptions required for the longer lifespan of taller species, which results in exposure to long-term infections and a myriad of stress sources. Importantly, *P. trichocarpa* had not been included in our analysis due to a large amount of gene duplication events detected by our pre-processing pipeline. Therefore, CALANGO independently provides evidence supporting the association of genes fulfilling these biological roles and the emergence of taller species.

Taller plants have lower rates of molecular evolution, presumably due to longer generation times and slower long-term rates of mitosis in their apical meristems. This fact is a concern for the long-term viability of these organisms and, consequently, for the large number of ecosystems where they play critical roles meristems (Lanfear, et al., 2013).

Self-Incompatibility (SI) systems are non-homologous molecular mechanisms that prevent inbreeding and promotes outcrossing in flowering plants (Durand, et al., 2020). Three of the domains associated with maximum height in angiosperms are components of the most well-characterized SI system (Figs 3C, “PF00954 – *S-locus glycoprotein domain*”; Supplementary Figure 6A-B; Supplementary Table 4, sheet “associated_domains”). The SI system found by CALANGO (from now on referred simply as “SI”) has been described in exquisite molecular details in the Brassicaceae family, even though it is widely distributed in flowering plants, and is an archetypal example of natural (balancing) selection maintaining genetic variation over long evolutionary times through inbreeding avoidance and rare-allele advantage (Durand, et al., 2020).

The SI system is a mating barrier controlled by a single highly polymorphic locus (S-locus), which codes for two genes in closely-linked genes: 1) the S-locus receptor kinase (SRK), a glycoprotein with kinase activity that allows stigma cells to discriminate between pollen from the same organism or from genetically related individuals (all four domains are located in these genes); 2) the S-locus cysteine-rich protein (SCR), expressed in pollen coat and the ligand of SRK (Nasrallah and Nasrallah, 2014). In the case of self-fertilization, the SCR protein in pollen is structurally complementary to the SRK protein found in the same S locus haplotype, activating a signaling pathway that inhibits pollen tube development. Not surprisingly, the S-locus is highly polymorphic, and has been extensively characterized in several populations across multiple species and in distinct ecological contexts (Durand, et al., 2020).

We found 37 copies of PF00954 – the signature domain of SRK genes – in *Arabidopsis thaliana* (maximum height of 0.30 meters, the eighth smallest plant in our dataset), while *Eucalyptus grandis* (maximum height of 55 meters, the tallest plant in our dataset) has 210 copies of this domain (a 5.68-fold increase). The remaining three components of the SI system have similar expansion profiles (Supplementary Figure 4, Supplementary Table 4, sheet “associated_domains”).

Our findings describe a previously unknown landscape of the total genetic variability in the SI system, as the combinatorial possibilities of multiple loci coding for components of the SI mechanism, each locus being itself potentially highly variable, greatly expands the total number of alleles hosted in each genome and, consequently, the fraction of possible incompatible individuals in the populations of taller plants. In this scenario, successful fertilization events would have a greater chance of outcrossing in species that can potentially host a greater number of S-locus haplotypes, therefore allowing taller plants to increase their evolutionary rates through cross-pollination with unrelated individuals.

CALANGO revealed a previously unknown molecular mechanism that can promote outcrossing in taller species with longer generation times and may counterbalance their lower evolutionary rates. This observation may have profound consequences for our understanding of the evolutionary future of taller plants and the critical roles they play in their ecosystems and in human economic activities, as the ability of a species to adapt to changing environments fundamentally depends on their underlying mutation rates (Willi and Hoffmann, 2009).

### Comparison of CALANGO with conceptually similar software

As recently reviewed by Lázló *et al*., several tools are already available to search for associations between the patterns of occurrence of homologous genomic components (mostly secondary losses of homologous regions in specific lineages) and a binary phenotype of interest (Nagy, et al., 2020). One important distinction between CALANGO and these tools is the class of phenotypic data accepted by them. Most of the methods for searching gene losses associated with categorical phenotypes (presence/absence) which, while useful to describe several types of biological variation, cannot be used to survey quantitative phenotypic data without the usage of *ad hoc* thresholds to define classes. Our tool, therefore, considerably expands the strategies currently available to search for associations between genetic and phenotypic data across species.

Furthermore, despite being successful in detecting gene losses associated with the emergence of complex phenotypes, none of these methods incorporate current genomic knowledge at the function level as provided by GO annotation. Instead, they exclusively evaluate the associations between sets of homologous elements across genomes and phenotypic data. GO terms have been shown to capture patterns of functional convergence and to provide a deeper biological comprehension of the genomic evolution of complex phenotypes, such as parasitism and sociality, and can provide a functional landscape for comparative genomics at the molecular function level (International Helminth Genomes, 2019; Tong, et al., 2020).

By dissociating the genomic components from their functional annotation, CALANGO provides flexibility to survey the distribution of several classes of genomic components, such as protein domains or entire genes. Furthermore, as we demonstrated, distinct annotation schemas allow both the emulation of classic comparative genomics analysis (by using an homology-based annotation dictionary) and a pathway- or function-based comparison (by using a GO-based dictionary), a feature that, to the best of our knowledge, is not present in any existing tools. As illustrated in our case studies, the combination of both strategies delivers a richer, more biologically meaningful interpretation of the results, including the detection of functional molecular convergences which could not be discovered using homology-based annotation.

Also in contrast with virtually all the software reviewed by Lázló, which mostly provide text files as main output, CALANGO produces a rich set of dynamical HTML5 result files containing several statistics and other useful quantities, together with their visual representations. For advanced users, CALANGO provides all results as a list of standard R objects, therefore allowing easy integration with other computational pipelines. The installation procedure for our tool is straightforward, as it is available as a fully operational R package.

## DISCUSSION

The post-genomic era has brought a plethora of high-quality sequenced genomes, ranging from previously underrepresented early-branching lineages of cellular organisms to thousands of genomes of a single bacterial species. In contrast to this abundance of genomic data, there are currently no standardized methods in computational statistics for extracting genomic properties associated with a quantitative genotype or phenotype of interest across genomes. CALANGO addresses this gap in the comparative genomics field by integrating phylogenetic, genomic, annotation and phenotypic data together in order to perform this task.

Our two case studies comprise datasets that are highly contrasting in terms of evolutionary time, taxonomic diversity and the quantitative phenotype/genotype under analysis. The first evaluated the biological roles associated with the change of a complex genotype (the density of prophages) in a single bacterial species. We found, as expected, a considerable association with genes of viral origin. By removing these genes and blocking their effect from the analysis, this case study allowed us to demonstrate how CALANGO can support the investigation of causal associations. We also observed several unknown associations at the function level that point to a much richer scenario of the biological interaction between bacteria, their prophages and other classes of mobile elements. We emphasize how the horizontal acquisition of adaptive genes, such as virulence factors and stress response genes, may allow bacteria to thrive in distinct environments.

The second case study detected domain expansions associated with a complex phenotype (maximum height) in the flowering plants, a major group of multicellular eukaryotes. Tall plants have morphological and physiological adaptions to the challenges of growing vertically (Falster and Westoby, 2003), and concomitantly harbor several advantages in dispersal and establishment success rates (Mashau, et al., 2021). Our case study described several mechanisms that greatly improve our understanding of the genomic regions and molecular mechanisms associated with emergence and maintenance of this complex phenotype. Of special interest, we described a previously unknown reproductive strategy that may allow taller plants the increase their genetical diversity through the independent expansion of the S-locus and the allocation of more resources to cross-pollination, a fact with long-reaching consequences for fields as diverse as biotechnology, agriculture and conservation biology. More importantly, our tool also indicated several testable hypotheses in both case studies, indicating how it can be used to prioritize downstream targets for experimental characterization.

CALANGO represents a considerable step towards the establishment of an annotation-based, phylogeny-aware comparative genomics framework to survey genomic data beyond sequence level, and to search for associations between quantitative variables across lineages sharing a common ancestor and the multiple layers of biological knowledge coded in their genomes.

## ABBREVIATIONS

QVAL: quantitative values across lineages,
DUF: domain of unknown function,
SRK: S-locus receptor kinase,
SCR: S-locus cysteine-rich protein.

## DATA AVAILABILITY

All processed data needed to fully reproduce the two case studies (genome annotation files, phylogenetic tree, phenotypic information and CALANGO configuration files) are available at https://labpackages.github.io/CALANGO/. All raw data used in case studies (genome IDs and sources for phenotypic data) are available as supplementary tables.

## SUPPLEMENTARY DATA

Supplementary Data are available in the online version of this article.

## ACKNOWLEDGEMENT

We would like to thank Prof. Daniel dos Santos Mansur and Prof. Glória Regina Franco for the insightful discussions during the elaboration of this research. We also would like to thank the, for the financial support for this research and publication fees.

## REFERENCES

Arndt, D., et al. PHASTER: a better, faster version of the PHAST phage search tool. Nucleic acids research 2016;44(W1):W16–21.

Bertozzi Silva, J., Storms, Z. and Sauvageau, D. Host receptors for bacteriophage adsorption. FEMS Microbiol Lett 2016;363(4).

Buchfink, B., Xie, C. and Huson, D.H. Fast and sensitive protein alignment using DIAMOND. Nat Methods 2015;12(1):59–60.

Chaudhuri, R.R. and Henderson, I.R. The evolution of the Escherichia coli phylogeny. Infect Genet Evol 2012;12(2):214–226.

Cornwell, W. and Nakagawa, S. Phylogenetic comparative methods. Curr Biol 2017;27(9):R333–R336.

Correa, A.M.S., et al. Revisiting the rules of life for viruses of microorganisms. Nat Rev Microbiol 2021.

Durand, E., et al. Evolution of self-incompatibility in the Brassicaceae: Lessons from a textbook example of natural selection. Evol Appl 2020;13(6):1279–1297.

Edgar, R.C. MUSCLE: a multiple sequence alignment method with reduced time and space complexity. BMC bioinformatics 2004;5:113.

Ehrbar, K. and Hardt, W.D. Bacteriophage-encoded type III effectors in Salmonella enterica subspecies 1 serovar Typhimurium. Infect Genet Evol 2005;5(1):1–9.

El-Gebali, S., et al. The Pfam protein families database in 2019. Nucleic acids research 2019;47(D1):D427–D432.

Falster, D.S. and Westoby, M. Plant height and evolutionary games. Trends in ecology & evolution 2003;18(7):337–343.

Felsenstein, J. Phylogenies and the Comparative Method The American Naturalist 1985;125(1):15.

Fischer, S., et al. Using OrthoMCL to assign proteins to OrthoMCL-DB groups or to cluster proteomes into new ortholog groups. Curr Protoc Bioinformatics 2011;Chapter 6:Unit 6 12 11-19.

Goodstein, D.M., et al. Phytozome: a comparative platform for green plant genomics. Nucleic acids research 2012;40(Database issue):D1178–1186.

Groth, P., et al. PhenomicDB: a new cross-species genotype/phenotype resource. Nucleic acids research 2007;35(Database issue):D696–699.

Hung, J.H., et al. Gene set enrichment analysis: performance evaluation and usage guidelines. Brief Bioinform 2012;13(3):281–291.

International Helminth Genomes, C. Comparative genomics of the major parasitic worms. Nat Genet 2019;51(1):163–174.

Jones, P., et al. InterProScan 5: genome-scale protein function classification. Bioinformatics 2014;30(9):1236–1240.

Kumar, S., et al. MEGA X: Molecular Evolutionary Genetics Analysis across Computing Platforms. Molecular biology and evolution 2018;35(6):1547–1549.

Kumar, S., et al. TimeTree: A Resource for Timelines, Timetrees, and Divergence Times. Molecular biology and evolution 2017;34(7):1812–1819.

Lanfear, R., et al. Taller plants have lower rates of molecular evolution. Nat Commun 2013;4:1879.

Lee, H., et al. Rate and molecular spectrum of spontaneous mutations in the bacterium Escherichia coli as determined by whole-genome sequencing. Proceedings of the National Academy of Sciences of the United States of America 2012;109(41):E2774–2783.

Li, W. and Godzik, A. Cd-hit: a fast program for clustering and comparing large sets of protein or nucleotide sequences. Bioinformatics 2006;22(13):1658–1659.

Liolios, K., et al. The Genomes On Line Database (GOLD) in 2009: status of genomic and metagenomic projects and their associated metadata. Nucleic acids research 2010;38(Database issue):D346–354.

Mashau, A.C., et al. Plant height and lifespan predict range size in southern African grasses. Journal of Biogeography 2021;48(12):3047–3059.

Moles, A.T., et al. Global patterns in plant height. Journal of Ecology 2009;97(5):923–932.

Mukherjee, S., et al. Genomes OnLine Database (GOLD) v.6: data updates and feature enhancements. Nucleic acids research 2017;45(D1):D446–D456.

Nagy, L.G., et al. Novel phylogenetic methods are needed for understanding gene function in the era of mega-scale genome sequencing. Nucleic acids research 2020.

Nasrallah, J.B. and Nasrallah, M.E. S-locus receptor kinase signalling. Biochem Soc Trans 2014;42(2):313–319.

O’Leary, N.A., et al. Reference sequence (RefSeq) database at NCBI: current status, taxonomic expansion, and functional annotation. Nucleic acids research 2016;44(D1):D733–745.

Pang, S., et al. GPU MrBayes V3.1: MrBayes on Graphics Processing Units for Protein Sequence Data. Molecular biology and evolution 2015;32(9):2496–2497.

Park, B.S. and Lee, J.O. Recognition of lipopolysaccharide pattern by TLR4 complexes. Exp Mol Med 2013;45:e66.

Peiffer, J.A., et al. The genetic architecture of maize height. Genetics 2014;196(4):1337–1356.

Quinlan, A.R. BEDTools: The Swiss-Army Tool for Genome Feature Analysis. Curr Protoc Bioinformatics 2014;47:11 12 11–34.

Rambaut, A., et al. Posterior Summarization in Bayesian Phylogenetics Using Tracer 1.7. Syst Biol 2018;67(5):901–904.

Ren, R., et al. Widespread Whole Genome Duplications Contribute to Genome Complexity and Species Diversity in Angiosperms. Mol Plant 2018;11(3):414–428.

Revell, L.J. phytools: An R package for phylogenetic comparative biology (and other things). Methods Ecol. Evol. 2012;3:7.

Steyert, S.R. and Kaper, J.B. Contribution of urease to colonization by Shiga toxin-producing Escherichia coli. Infect Immun 2012;80(8):2589–2600.

Tong, C., et al. Comparative Genomics Identifies Putative Signatures of Sociality in Spiders. Genome Biol Evol 2020;12(3):122–133.

Touchon, M., et al. Organised genome dynamics in the Escherichia coli species results in highly diverse adaptive paths. PLoS Genet 2009;5(1):e1000344.

Tuskan, G.A., et al. The genome of black cottonwood, Populus trichocarpa (Torr. & Gray). Science (New York, N.Y 2006;313(5793):1596–1604.

Vaidya G. L.D., Meier R. SequenceMatrix: concatenation software for the fast assembly of multi-gene datasets with character set and codon information. Cladistics 2011;27(2):9.

Vogel, C. and Chothia, C. Protein family expansions and biological complexity. PLoS Comput Biol 2006;2(5):e48.

Wang, X., et al. Cryptic prophages help bacteria cope with adverse environments. Nat Commun 2010;1:147.

Waterhouse, R.M., et al. BUSCO Applications from Quality Assessments to Gene Prediction and Phylogenomics. Molecular biology and evolution 2018;35(3):543–548.

Willi, Y. and Hoffmann, A.A. Demographic factors and genetic variation influence population persistence under environmental change. J Evol Biol 2009;22(1):124–133.

Wolf, A.J. and Underhill, D.M. Peptidoglycan recognition by the innate immune system. Nat Rev Immunol 2018;18(4):243–254.

Zu, P. and Schiestl, F.P. The effects of becoming taller: direct and pleiotropic effects of artificial selection on plant height in Brassica rapa. Plant J 2017;89(5):1009–1019.

