## Supplementary File 1 for "CALANGO: a phylogeny-aware comparative genomics tool for discovering quantitative genotype-phenotype associations across species"

**SUPPLEMENTARY RESULTS**

**Integrated bacteriophages in pathogenic and non-pathogenic *E. coli***

We selected 80 high-quality, gapless *E. coli* genomes (chromosomes and plasmids) that could be unambiguously classified as either pathogenic (51) or non-pathogenic (29) lineages, excluding laboratory strains (Supplementary Table 1). We predicted the numbers and location of prophages in *E. coli* chromosomes using PHASTER (Arndt et al., 2016) and found 729 prophages, with counts varying from one to 22 integrated viruses per *E. coli* genome (Figure 2A).

We found pathogenic lineages to have a significantly higher count of prophages when compared with non-pathogenic ones (Figure 2A, “Prophage count”). We also observed pathogenic lineages to possess genome sizes significantly larger than non-pathogenic lineages (Figure 2A, “Genome length”). To exclude the possibility that pathogenic *E. coli* have more integrated prophages due to their larger genomes we proceeded by computing prophage densities for each genome (the number of prophages divided by genome length), which was also found to be significantly higher in pathogenic lineages (Figure 2B, “Prophage density”). We concluded that pathogenic *E. coli* have an increased prophage occurrence in their genomes even after accounting for genome size and phylogeny as a possible source of bias. We proceeded by using phage density as a QVAL to survey the *E. coli* dataset for biological functions associated with prophage occurrence.

**Comparison of functional and homology-based annotation schemas**

We investigated whether a GO-based annotation effectively integrates information from non-homologous domains that fulfill the same biological role by evaluating the prevalence of annotation terms, defined as the percentage of genomes where they were observed. This is a useful statistic to evaluate whether a GO-based annotation, which is expected to capture non-homologous functional similarities, is more observed across genomes when compared with a homology-based annotation schema. The sum of annotation term occurrence in all genomes also provides a proxy for the overall abundance of annotation terms in distinct annotation schemas. If a GO-based annotation of a set of genomic components integrates the information from non-homologous coding regions that fulfill the same biological roles, we expect it to have both a greater prevalence and a greater sum value than a Pfam-based annotation of the same set of components. Additionally, we also expect these initial differences in the occurrence of annotation terms to be reflected in the annotation terms found to be significantly associated with prophage density.

The distribution of prevalence values in all the 3729 annotation terms from *domain2Pfam* analysis observed in at least one genome revealed a highly skewed distribution, where most annotation terms are observed in the vast majority of genomes (Supplementary Figure 1C, “Prevalence” chart, “domain2Pfam all”). We found that 65.7% of the Pfam IDs (2450 out of 3729) are observed in more than 90% of the *E. coli* proteomes. A minor, second distribution peak is observed in Pfam IDs with the smallest prevalence values, comprising 11.6% of the Pfam IDs with prevalence < 0.1 (432 out of 3729 Pfam IDs).

The distribution of GO terms used to annotate the same set of genomic components (domain2GO experiment) has an even more biased distribution regarding terms with high prevalence values, where 88.8% (1768 out of 1992 terms found annotating at least one Pfam entry) have prevalence values greater than 0.9 (Supplementary Figure 1C, “Prevalence” chart, “domain2GO­_all”). Furthermore, only 2.3% of the GO terms have prevalence values smaller than 0.1 (46 out of 1992). This indicates that, even though the majority of terms from both functional- and homology-based annotation schemas are observed in most genomes, the terms from the functional-based schema are significantly more prevalent than those from a homology-based one (Wilcoxon test, p-value < 2.22e-16). The higher prevalence observed in our GO-based annotation suggests this annotation schema is indeed capable of better representing common biological themes shared across genomes than a homology-based one.

The higher values of prevalence observed in annotation terms from *domain2GO* appear to be reflected in significantly associated terms from this annotation schema (Supplementary Figure 1C, “Prevalence” chart, “domain2GO_all” and “domain2GO_sig”), which were also found to be highly biased towards high prevalence values (80.8%, 177 out of 219). The observed difference in the median prevalence from all terms from *domain2GO* annotation and the significant terms from the same annotation was not found to be statistically significant (Wilcoxon test, p-value = 0.057).

In contrast to *domain2GO* annotation, the significant annotation terms from the *domain2Pfam* experiment have much lower prevalence values than the distribution observed in all *domain2Pfam* annotation terms (Supplementary Figure 1C, “Prevalence” chart, “domain2Pfam_all” and “domain2Pfam_sig”). Only 25,7% (59 out of 230) of the significant annotation terms have prevalence values greater than 0.9. We also found the median prevalence value for the associated terms to be significantly lower than both the one observed for all annotation terms from the *domain2Pfam* experiment and the significant terms from *domain2GO* experiment (Wilcoxon test, p-values < 2.22e-16).

We further evaluated annotation term occurrence in homology- and functional-based annotation schemas by computing the sum of occurrences of each annotation term in each annotation schema (Supplementary Figure 1C, “Sum” chart). We initially observed that the homologous regions have a significantly smaller sum of occurrences than GO terms (medians of 80 and 227, respectively; Wilcoxon test, p-value < 2.22e-16). The distribution of significant annotation terms from homology-based annotations schema was also found to be more similar to the distribution of sums of all *domain2Pfam* terms (medians of 108.5 and 80, respectively; Wilcoxon test, p-value = 3.6e-04) than the significant associations of functional-based annotations when compared with the sum of term occurrences for *domain2GO* experiment (medians of 1125 and 227, respectively for associated GO IDs and all GO terms; Wilcoxon test, p-value < 2.22e-16). Furthermore, associated GO terms have significantly higher sum values than associated Pfam IDs (medians of 1125 and 108.5, respectively; Wilcoxon test, p-value < 2.22e-16).

The scenario where annotation terms from a GO-based annotation schema are both more prevalent and more abundant than terms from the Pfam-based annotation is compatible with the several classes of genes non-homologous genes of viral origin that play complementary roles in the viral life cycle found to be associated with prophage densities. These observations are also compatible with the integration of biological knowledge at the function level, where non-homologous Pfam domains observed in distinct genomes are annotated using the same GO term, therefore increasing both the prevalence and the abundance of GO terms in comparison to Pfam IDs.

**Evaluating annotation term frequencies and counts**

When evaluating annotation terms associated with QVAL, CALANGO allows users to use either raw count data of annotation terms or to normalize these values to account for biologically meaningful differences across genomes. This normalization procedure may be interesting when evaluating genomes that have, for instance, considerable variation in their number of protein-coding genes, which may cause annotation term counts to be more abundant in genomes with larger numbers of genes.

On the other hand, normalization using a common factor induces dependencies in the relative frequencies for each annotation term, as the resulting values must add to one (Supplementary Figure 2A). This causes a mathematical artifact that can produce spurious significant associations of frequency of terms having constant or near-constant count values, due to relative frequencies of other terms whose counts are associated with QVAL.

Supplementary Figure 2A illustrates five hypothetical genomes with two annotation terms, one occurring once in each of them (e.g., a universal 1-1 ortholog, represented as an orange box), and another having a variable count occurrence that is associated with a QVAL vector (blue box). When computing associations from count data, we only observe the blue term to be associated with QVAL (Supplementary Figure 2A, “Blue count” and “Orange count” plots), as expected.

When examining relative frequencies, however, we notice the emergence of a spurious association of orange frequencies with the QVAL, in addition to the expected association of blue terms (Supplementary figure 2A, “Blue freq” and “Orange freq” plots). This results from the dependency structure between the blue terms and the QVAL “contaminating” the QVAL-independent orange counts via the normalization denominator, which results in a negative correlation between the orange and blue frequencies (Supplementary Figure 2A, “Blue & Orange freq” plot).

In this work we initially used CALANGO to survey both annotation term counts and relative frequencies for possible associations with prophage densities (Supplementary Table 2, Supplementary Figure 2B). Although the majority of positively associated terms were observed to be the same when considering either count or relative frequencies, we found most of the negatively associated terms to be detected only when considering relative frequencies (Supplementary Figure 2B).

Furthermore, manual inspection of the annotation terms detected in the second case found a considerable fraction of them to correspond to core housekeeping cellular processes (Supplementary Table 2), further suggesting that their counts are likely to be near constant across genomes, and that the negative associations found in frequency data is likely to be an artifact caused by the issue highlighted earlier. Taken together, these experiments demonstrate that relative frequencies are not always the most adequate representation of annotation term occurrence, which motivated our decision to proceed with count data alone for the *E. coli* dataset.

**Homologous genes and biological roles associated with prophage density in *E. coli***

Figure 2 shows an annotated version of the heatmap produced by CALANGO using *domain2Pfam*, representing all Pfam domain IDs associated with prophage density. This figure demonstrates how our tool integrates phylogenetic, phenotypic, and annotation data to allow downstream exploratory analysis. Species clustering is done using the phylogenetic tree, together with visualization of user-defined groups (pathogenic and non-pathogenic *E. coli* lineages, in our case), allowing the detection of interesting distribution patterns of annotation terms while considering, in our case, phylogenetic and phenotypic/ecological information. Pathogenic and non-pathogenic lineages are distributed with no clear broader grouping pattern, suggesting that both phenotypes emerged and/or were lost several times during the evolution of these lineages. Two pathogenic *E. coli* groups have the highest count of most Pfam domains (Figure 2, red arrows). These lineages comprise enterohemorrhagic (EHEC) and enteropathogenic (EPEC) pathotypes, including all O157:H7 lineages (larger group), a Shiga-like toxin-producing serotype, and an important source of foodborne disease (Steyert and Kaper, 2012).

We found the domain clustering to comprise mostly two major functional classes: groups of non-homologous regions of viral origin that play several roles in the viral life cycle, such as integrases, structural proteins, lysozymes, and DNA metabolism enzymes, together with three main classes of virulence factors known to play important roles in EHEC and EPEC disease etiology: Type III secretion systems, urease, adhesins, hemolysins, NFkB-degrading protease, attaching and effacing virulence factors, and Shiga-like toxin (Jarvis et al., 1995; Steyert and Kaper, 2012; Wu et al., 2010). The two groups of pathogenic *E. coli* with the highest count of associated Pfam IDs also contain most of the genomes where the virulence factor clusters were observed.

**Positive association: virulence factors and insertion elements**

The second most abundant category of protein domains found to be associated with prophage density were several classes of virulence factors. Interestingly, we found that 44 out of the 58 (75,9%) of domains annotated by us as virulence factors are still significantly associated with prophage density even after the removal of viral genes, suggesting that a considerable fraction of such genomic elements is not located within predicted prophage genomes (Supplementary Table 2, sheet “virulence_factors”, Supplementary File 1, section “Annotation terms associated with prophage density after removal of genes of viral origin”). In fact, from the 4379 domains coding for virulence factors, 3122 (71.3%) are located outside detectable regions of viral origin.

We observed that 7 out of the 14 virulence factor domains not associated with prophage density after the removal of the genes of viral origin are components of pathogenicity mechanisms long known to be horizontally transferred by bacteriophage integration, such as Shiga-like toxins and effectors of Type III secretion system (Ehrbar and Hardt, 2005; Steyert and Kaper, 2012) (*e.g.* all 22 copies of domain PF02258 - *Shiga-like toxin beta subunit*, Supplementary Table 2). This provides additional evidence that our strategy to remove the effect of genes of viral origin was successful.

The unexpected observation that most virulence factors are located outside predicted prophages and remain associated with the QVAL after the removal of genes of viral origin suggests a more complex relationship between virulence factors and prophage occurrence. One possibility is that the presence of these domains is a consequence of previous bacteriophage integration events that, over time, resulted in 1) prophage degeneration by the loss of genes needed for viral replication, as a result of selective pressure for smaller bacterial genomes with lower replication times; and 2) the maintenance of genes contributing to a pathogenicity phenotype due to a fitness increase provided by such genes in pathogenic *E. coli* lineages (Ramisetty and Sudhakari, 2019) - a scenario compatible with the association of domain PF06316 after the removal of viral genes (Supplementary Figure 3E). A non-excluding alternative hypothesis is that some of these virulence domains may have arrived in bacterial genomes by non-phage-mediated horizontal transfer events and, together with virulence factors acquired through bacteriophage integration, contribute to a pathogenic phenotype. More complex hypotheses are also reasonable, such as the occurrence of synergism between the presence of a prophage and the external virulence factor, resulting in increased fitness to the bacteria when both are present.

We found five domains coding for insertion elements to be positively associated with prophage density, even after the removal of genes of viral origin (Supplementary Table 2, sheet “domain2PfamCountLessPhages”, PF13007 - *Transposase C of IS166 homeodomain*). Despite having been initially described as classic deleterious genomic parasites, insertion elements have been increasingly documented as important agents of bacterial genome evolution by providing the horizontal acquisition of genomic components that confer adaptation to new niches, including virulence factors, and, interestingly, may provide resistance to bacteriophage infection (Vandecraen et al., 2017).

Although there is plenty of evidence that bacteriophage integration contributes to pathogenic phenotypes in bacterial species through horizontal gene transfer of virulence factors [32, 36], we found no previous studies demonstrating a significant association between prophage density in bacterial genomes and the occurrence of virulence factors, while correcting for possible confounders and known biases, such as phylogeny and genome quality. Furthermore, our results demonstrate that the majority of virulence factors remain associated with prophage density even after the removal of genes of viral origin, therefore excluding the contribution of all known genes of viral origin for these associations. These unforeseen results showcase how CALANGO can be used to enable controlled *in silico* experiments for the investigation of biologically meaningful questions.

**Positive association: other biological roles**

Besides protein domains associated with phage biology, virulence factors, and insertion elements, CALANGO detected 36 additional domains positively associated with prophage density that fulfill several biological roles. Remarkably, these include four domains that code for anti-viral components of bacterial immunity, such as putative components of the restriction-modification and CRISPR-Cas systems (e.g. PF13588 - *Type I restriction enzyme R protein N terminus (HSDR_N)*, Supplementary Figure 3G). It may be tempting at this point to assume that integrated prophages constitute a selective pressure for the occurrence and maintenance of such mechanisms. However, these four domains are not associated with prophage density after removal of genes of viral origin, therefore suggesting that at least a fraction of them occur within prophage genomes. We hypothesize these could represent viral genes related to the hijacking of bacterial immunity to prevent, for instance, infection by competing viruses, but further studies are needed for confirmation. Furthermore, some components of anti-viral mechanisms were also found to be negatively associated with prophage density, as we shall see, suggesting a more complex interplay between integrated prophages and the occurrence of anti-viral mechanisms (Supplementary File 1, section “Distinct anti-viral mechanisms are positively and negatively associated with prophage density”).

We also found 10 positively associated DUFs that may either comprise domains of viral origin playing roles in viral life cycles, or genes performing other biological functions found to be associated with prophage density, such as previously unknown virulence factors (Supplementary Table 2). Three out of the 10 DUFs remain significantly associated with the QVAL even after the removal of all predicted prophage genes, indicating that at least a fraction of them is located outside detectable integrated viral genomes. This property is shared with several of the known virulence factors we observed, making these domains interesting targets for downstream functional evaluation.

**Biological roles negatively associated with prophage density**

A total of 23 domains were found to have significant negative associations with prophage density (Figure 2), and 20 of them are associated after the removal of genes of viral origin. Most of these domains suggest a scenario where *E. coli* lineages with fewer integrated prophages, which tend to be non-pathogenic, have a set of genes enabling a greater diversity of lifestyles at several levels. At the molecular/metabolic level, we found domains coding for metabolic functions (e.g. PF00881 − *Nitroreductase family*, Supplementary Figure 3L) and transmembrane transporters involved in the uptake of ions and organic molecules (e.g. PF00950 - *ABC 3 transport family*, Supplementary Figure 3K).

We found two domains representing distinct facets of bacterial diversity at the population level, the first coding for a putative glycosyl hydrolase described with a known role in biofilm formation (PF14883 - *Hypothetical glycosyl hydrolase family 13* (Nishiyama et al., 2013). Bacteriophages can modulate biofilm formation in several bacterial species, and certain phages are efficient biofilm destroyers (Fernandez et al., 2018). However, inside biofilms, susceptible bacterial populations could be protected from phage infections by their resistant counterparts (Simmons et al., 2020). In this case, the presence of the PF14883 domain in the bacterium could be a protective factor against phages, and explain the negative association detected by CALANGO. The second domain (PF17508 - *Microcin V bacteriocin*, Supplementary Figure 3J) codes for a bacteriocin, an umbrella term used to describe protein toxins produced by bacteria that inhibit the growth of closely related species (Cotter et al., 2013). The death of related competitors could be a protective measure against phages, allowing a bacteriocin-producing population to keep in check potential viral spreaders in its surroundings during kill-the-winner dynamics (Maslov and Sneppen, 2017).

Phage-bacteria interactions inside a metazoan have been shown to differ from those happening in the environment or in laboratory conditions. At the metazoan-environment interface (mucosal layers), the pressure from phages might be greater than elsewhere, considering the impact of enriched mucosal-associated phages (Barr et al., 2013). Also, the mucosal layer could influence lysis-lysogeny decisions, and prophage-containing bacteria have been speculated to be protected in this environment by superinfection exclusion (Silveira and Rohwer, 2016). This correlates well to CALANGO findings that pathogenic *E. coli* tends to have more associated prophages, as the inserted viral genes could provide protection against phages colonizing the metazoans during the bacterial invasion process. This situation could also explain why some virulence factors coded by the bacteria are positively associated with prophages: the prophage-mediated protection could add up to the virulence gain, synergistically increasing the bacterial fitness in pathogenic situations.

We also observed a negative association of a component of an anti-viral mechanism, even after removing genes of viral origin (PF04313 - *Type I restriction enzyme R protein N terminus (HSDR_N)*), suggesting this domain may play some role in preventing viral infections in the lineages with a smaller prophage occurrence (Supplementary File 1, section “Anti-viral mechanisms are positively and negatively associated with prophage density”).

Three interesting examples are the negative associations for two glycosyltransferase domains (PF08437 and PF01501) that are components of the lipopolysaccharide (LPS) biosynthesis pathway, and for one peptidoglycan binding domain (PF01471) found in genes involved in cell wall processes (Figure 2F, Supplementary Figure 3I). LPS and peptidoglycan are components of cell wall and have roles both as major players of *E. coli* virulence, but also as receptors of several bacteriophages during adsorption.

LPS are a major component of cell walls of gram-negative bacteria and, together with peptidoglycan, are also receptors of several bacteriophages during adsorption to the bacterial cell surface when starting the infection process (Bertozzi Silva et al., 2016). Therefore, the decrease of these domains may be a consequence of the selective pressure caused by bacteriophage infection, resulting in surface modifications leading to the loss or change of molecules used by the phages as viral receptors. Acquisition of phage resistance through surface modification is detrimental to bacterial virulence for different pathogenic bacterial species (Gordillo Altamirano et al., 2021; Leon and Bastias, 2015).

However, LPS molecules have also been long recognized as major activators of the innate branch of the vertebrate immune system (Park and Lee, 2013). Consequently, the decrease of these domains may also represent evidence of the selective pressures of vertebrate immune systems on pathogenic bacteria, resulting in a decrease in *E. coli* immunogenicity through the loss of bacterial cell wall components. It is possible that both effects may be happening concomitantly, with the selective pressure induced by prophage infection resulting in the decrease of genes coding for components of LPS and other components of the cell wall, but also causing an exaptation scenario where this selection enables pathogenic *E. coli* to better adapt to its niche by partially evading host immune system. This illustrates another powerful aspect of our software, namely its ability to elicit relevant testable hypotheses from the data. In the particular case of this study, it would be possible to evaluate the relative fitness contribution of these domains through targeted knockout in distinct *E. coli* lineages, followed by controlled experiments investigating vertebrate immune system evasion or prophage infection rates.

**Anti-viral mechanisms are positively and negatively associated with prophage density**

We found five Pfam IDs associated with prophage density that are also components of bacterial immune systems to prevent viral infections, with four positive associations and one negative. Among the positive associations we found two DNA methylases (PF05063 - *MT-A70* and PF01555 - *DNA methylase*), one DNA-binding domain found in CRISPR negative transcriptional regulators (PF13412 - *Winged helix-turn-helix DNA-binding*) (Pawluk et al., 2014) and a restriction component of type I restriction-modification systems (PF13588 - *Type I restriction enzyme R protein N terminus (HSDR_N)*). All positive associations are not observed after the removal of genes of viral origin, indicating that a considerable fraction of these domains is found within prophages (Supplementary Table 2, sheet “domain2PfamCountLessPhages”).

Interestingly, both the DNA methylases and the negative transcriptional regulator may confer advantages for bacteriophages to evade bacterial immune systems. As for PF13588, it is worth noting that we also found a domain described as a component of restriction-modification mechanism to be negatively associated with prophage density (PF04313 - *Type I restriction enzyme R protein N terminus (HSDR_N)*), suggesting that some restriction systems may occupy distinct biological roles in extremes of phage density. A possible hypothesis is that some restriction-modification systems may be horizontally transferred by bacteriophage genomes and confer bacterial resistance to additional bacteriophage phage infections to avoid competition (Dedrick et al., 2017). As for the negatively associated restriction system, it is a component of bacterial genomes observed outside prophage regions that may provide a more general bacteriophage infection resistance in *E. coli* with few integrated prophages. Also, the loss of such systems in lineages with greater values of prophage density may suggest such loss may be advantageous for a parasitic lifestyle. We again highlight that CALANGO output produces hypotheses that are testable through genome edition of specific genomic components followed by relative fitness evaluation in controlled environments.

**GO annotation provides curation-level of biological knowledge**

The *domain2Pfam* analysis required extensive curation of domain functions to provide proper biological context to our findings. Furthermore, protein domain IDs represent homologous conserved regions within proteins and do not capture biological knowledge, such as functional similarities shared by non-homologous genomic components.

As CALANGO models genomic components independently from annotation terms, it is possible to objectively evaluate the influence of distinct annotation schemas used to annotate the same set of genomic components (Supplementary Figure 1B). At this point, we are interested in evaluating if GO annotation (*domain2GO*) would detect the same major biological themes observed during our manual curation of *domain2Pfam* results. Additionally, we evaluated if the integration of biological knowledge at the function level through GO annotation allows the detection of biological functions associated with prophage density that was not immediately discernible in our *domain2Pfam* analysis.

From the set of 1963 GO terms found to annotate at least five distinct protein domains as predicted by Pfam, CALANGO found 217 (11%) to be significantly associated with prophage density, with 195 positively correlated terms (correlations between 0.25 and 0.86) and 26 negatively associated (correlations between -0.27 and -0.54) (Supplementary Table 2, sheet “domain2GOCount”). As observed in the *domain2Pfam* analysis, we again found the majority of the positive associations to represent major aspects of bacteriophage biology and life cycle, including both general concepts and more specific components of the lytic and lysogenic cycles (e.g. GO:0019058 (*viral life cycle*) and GO:0019068 (*virion assembly*) (Figure 2G-H, Supplementary Figure 4 A-F, Supplementary Table 2, sheet “domain2GOCount”).

The protein domains annotated to these GOs comprise non-homologous sequences that play complementary roles in bacteriophage life cycles and their relationships with its bacterial host, such as in the several structural components of viral particles (e.g., portal proteins, head-to-tail joining proteins, tail proteins, head proteins, and capsids). Together, these results demonstrate how CALANGO integrates information from non-homologous sequences at the function level to automatically provide the same functional roles found in *domain2Pfam* analysis through manual curation and literature review.

CALANGO also detected associated GO terms representing general and specific aspects of *E. coli* pathogenicity to be the second-largest group of annotation terms positively associated with prophage density. The more general GO term GO:0009405 (pathogenesis) annotates several non-homologous virulence factors, such as cell invasion and adhesion, toxins, hemolysins, colicins and components of secretion systems, and provides further evidences of how CALANGO finds associations (Figure 2H). We also found other GO terms representing specific pathogenicity modules and mechanisms known to play important roles in distinct *E. coli* pathotypes, such as type III secretion and urease activity [46, 47] (Supplementary Figure 4G-J).

The 26 GO terms negatively associated with prophage density largely reflect the conclusions achieved at the domain-level analysis (*domain2Pfam*) after extensive manual curation (Supplementary Table 2, sheet “domain2GOCount”). We again found negative associations of GO terms reflecting a higher metabolic diversity (e.g.*,* GO:0043190 - *ATP-binding cassette (ABC) transporter complex*; GO:0072330 - *monocarboxylic acid biosynthetic process*) and also terms indicating the previous negative association between components of the LPS biosynthesis pathway (e.g., GO:0008918 - *lipopolysaccharide 3-alpha-galactosyltransferase activity*, Supplementary Figure 4K-N). Together, these results provide further evidence that, starting from the same set of genomic elements annotated to distinct dictionaries of biological roles, CALANGO can automatically detect biologically meaningful associations that are comparable to our manual curation of the results and equivalent to traditional comparative genomics analysis.

**Stress response genes are associated with prophage density in *E. coli***

We found 54 Pfam domains annotated as stress response mechanisms that play roles in several stress-related biological processes (Supplementary Table 2, sheet “stress_response_genes”). Approximately 30% these domains code for core biological functions observed mostly as single-copy universal orthologs, such as components of DNA repair pathways, oxidative stress pathways, transcription factors, chaperones and heat shock proteins. As they do not vary across genomes, they may not be the ones accounting for the association of GO:0006950 and prophage density.

The other 38 domains were observed in accessory proteins found in some *E. coli* lineages but not in others. Among them we observed components of restriction-modification systems, DNA repair pathways, colicins, toxin-antitoxin systems, Tellurite resistance, and transcription factors. Only two of these 38 domains were also detected in *domain2Pfam* experiment, indicating again that most of these Pfam IDs are not individually detectable as associated with prophage density but, when annotated to GO, the variation patterns of these genomic elements are aggregated in the function level and eventually contribute for the association to emerge. Additionally, a total of five (13.16%) of these domains are more represented in regions of viral origin than in host’s genomes, and 28 of them (73.7%) are found in at least one bacteriophage genome, indicating that the pool of stress response domains is present in both host and viral genomes.

Among the stress-response domains we also found some candidates for cellular responses to viral infections, such as the DNA repair pathways and restriction-modification mechanisms. It may be appealing to assume that an increase in genes fulfilling this biological role is the consequence of an adaptation process of host cells to the presence of integrated prophages. However, domains belonging to these categories were observed both in regions of viral origin and host’s genomes, providing additional evidence that anti-viral mechanisms carried out by bacteriophages may comprise an advantage for both hosts and integrated viruses by preventing competition through additional viral infection. We also found other stress response domains that may confer a fitness increase, such as virulence factors, warfare mechanisms and defense systems.

**Associations with prophage density after removal of genes of viral origin**

When searching for biological functions associated with prophage density in *E. coli* genomes, we know beforehand the location of all predicted prophages. Therefore, it is possible to remove all genes predicted as having viral origin (Supplementary File 1, section “Removal of genes of viral origin”) and objectively evaluate the effect of this procedure on associated annotation terms. Such genomes lacking genes located within prophage genomes were annotated using InterProScan (Jones et al., 2014), ordered according to their original prophage densities before the removal of viral genes and evaluated using CALANGO with the same criteria for significance.

For the *domain2Pfam* annotation schema after excluding genes of viral origin, we found 86 Pfam IDs still associated with prophage density (Supplementary Table 2, sheet “domain2PfamCountLessPhages”), 57 of which in common with the ones found in the original *domain2Pfam* experiment including viral genes (Supplementary Table 2, sheet “domain2PfamCount”). The 125 distinct Pfam IDs manually curated as of viral origin and associated with prophage density were found to occur 23,310 times across the *E. coli* proteomes. The removal of known genes of viral origin decreased the occurrence of these domains to 6,040, a reduction of 74%, therefore demonstrating we have been able to remove the majority of such domains. Furthermore, only two out of 125 domains were still found to be associated with prophage density in this experiment.

The first one is PF06316 (*Enterobacterial Ail/Lom protein*), a protein domain found virulence-related outer membrane protein family that is observed in bacteriophage genes, where it plays a role in lysogenic cycles (Pulkkinen and Miller, 1991), and also contributes to a pathogenicity phenotype in gram-negative bacteria by allowing both resistance to complement activity and the ability to adhere and invade host cells (Cirillo et al., 1996). Furthermore, this domain has 289 copies in bacterial genomes before the removal of genes of viral origin, but only 14 copies (4.84%) remain after the removal of such genes. These observations suggest a scenario of bacteriophage-mediated horizontal gene transfer followed by prophage degeneration and the eventual maintenance of virulence factors in pathogenic lineages as a consequence of fitness increase. The second Pfam domain is PF07799 (*Protein of unknown function (DUF1643)*), a DUF found in several proteins in Archaea, Bacteria and bacteriophages that remains to be characterized (Jones et al., 2014) and was observed in 11 copies in both experiments. Together, these data indicate that the mechanism that associates the annotation terms previously classified through manual curation as having roles in the viral life cycle and found to be associated with prophage density is indeed the presence of genes of viral origin, rather than by other potential mechanisms.

As for the set of 56 protein domains expected to contribute to a pathogenicity phenotype in *E. coli*, 42 of them (75%) are still significantly associated with prophage density after the removal of genes of viral origin (Supplementary Table 2, sheets domain2PfamCount” and “domain2PfamCountLessPhages”). Additionally, 27 and 39 of such domains have exactly the same number of occurrences or differ by one, respectively, when comparing the output of the *domain2Pfam* experiments with and without viral genes (Supplementary Table 2, sheet “virulelence_factors”). In contrast with the protein domains annotated as playing roles in the viral life cycle, the removal of viral genes did not alter the occurrence of the majority of virulence factors, as most of them are located outside predicted prophages.

We performed the same *in silico* procedure in *domain2GO* annotation schema to remove genes of viral origin and evaluate GO terms that remain associated with prophage density, again finding that the vast majority of annotation terms describing viral lifestyle functions not to be significantly associated with prophage densities once viral genes are removed (Supplementary Table 2, sheet “domain2GOCountLessPhages”). Most GO terms describing pathogenicity mechanisms, on the other hand, are also observed in *E. coli* genomes after removing the genes of viral origin, suggesting most of the domains annotated to these GO terms are located outside the regions of integrated prophages.

**Protein domains independently expanded in taller angiosperms**

It is worth noting that the domain with the domain with the smallest value of occurrence in our dataset and that is also associated with maximum height was observed 1108 times (PF11721, with 16 copies in *A. thaliana* and 62 copies in *E. grandis*, Supplementary Table 4, sheet “associated_domains”). Therefore, these domains comprise relatively large expansions in both absolute and relative terms, and therefore are not likely to be artifacts caused by the common pitfalls found in genome assembly and annotation, a particularly challenging field in plant genomics (Salzberg, 2019).

The Malectin domain (PF11721) was initially characterized in the model organism *Xenopus laevis*, where it monitors protein glycosylation in the endoplasmic reticulum (Schallus et al., 2008). However, the crystal structure of this domain in plants revealed the absence of critical amino acids for the interaction with diglucosidues that are present in the animal enzymes, suggesting distinct functional roles for proteins containing this domain in these lineages (Xiao et al., 2019). We found this domain to have 15 and 62 copies *A. thaliana* and *E. grandis*, respectively (4.13X increase) (Figure 3C, Supplementary Table 4, sheet “associated_domains”).

In *A. thaliana*, enzymes containing this domain have been functionally characterized as cell wall sensors that regulate development, reproduction and resistance to various stresses (Kumar et al., 2020). Interestingly, this domain has been previously reported to be greatly expanded in land plants when compared with other eukaryotes (Yang et al., 2021), and also expanded in the genome of *Populus trichocarpa*, a model organism for tree plant biology, when compared with *A. thaliana* (Kumar et al., 2019). Genes containing this domain are also upregulated in the developing wood tissue of *P. trichocarpa* and *E. grandis* (Kumar et al., 2019; Pinard et al., 2015), and the expansion of malectin-containing genes in *P. trichocarpa* appears to be a key player in the development of wood tissue in this species (Kumar et al., 2020).

We have not used *P. trichocarpa* in our analysis, as this species has been subject of a whole-duplication event and therefore failed to fulfill our genome quality metrics (Tuskan et al., 2006). Therefore, the data automatically produced by CALANGO independently strengthens the hypothesis that malectin-containing genes play a role in wood tissue development and are independently expanded in taller plants. This is yet another showcase of how CALANGO can be used to find potential causal relationships that can be surveyed through downstream experiments.

Ankyrin repeats (PF12796) mediate protein-protein interactions in a wide range of biological processes, and is one of the most common and phylogenetically diverse domain in public sequence databases, being observed in viruses, prokaryotes and eukaryotes, with the later comprising the vast majority of entries (Mosavi et al., 2004). We found 126 and 530 copies of PF12796 in *A. thaliana* and *E. grandis*, respectively (4.12x increase, Figure 3C). In *A. thaliana* proteins containing this domain play several roles in early embryogenesis and organ development (e.g. ANK6 - *ankyrin repeat protein 6*, KEG - *KEEP ON GOING*, EMB506 - *embryo defective 506*), even though a considerable fraction of these genes lacks functional characterization and are, therefore, interesting targets for future functional characterization.

Two domains (PF00560 and PF13855) comprise leucine-rich repeats, whose are frequently involved in protein-protein interaction processes. In the flowering plants, these domains are commonly found in Leucine-Rich Repeats Receptor-Like Kinases (LRR-RLKs), which is the case for the majority of *A. thaliana* genes (Supplementary Table 4, sheet “Arabidopsis_genes”). LRR-RLKs is one of the largest and most complex gene family in this species, playing roles in developmental pathways and immunity, perception of environmental conditions and stress response (Dufayard et al., 2017). We found 179 and 582 copies of domain PF00560 in the *A. thaliana* and *E. grandis* non-redundant proteomes, respectively (3.25x increase). As for PF13855, these proteomes have respectively 439 and 1295 copies of it (2.95x increase).

In *A. thaliana*, several of the genes containing these domains lack functional characterization, which is certainly true for an even greater fraction of *E. grandis* genes. The characterized genes in the thale cress that code for these domains are highly diverse in their functional aspects. We found this domain to occur in regulators of floral development (BAM1 - *BARELY ANY MERISTEM 1*, AT1G11130 - *STRUBBELIG*), modulators of controlled cell death (SERK5 - *somatic embryogenesis receptor-like kinase 5*), components of hormone signaling pathways (AT3G13380 - *BRI1-LIKE 3*), and agents of plant resistance to pathogens (AT4G26090 - *RESISTANT TO P. SYRINGAE 2*). The expansion of LRR-RLKs suggests that both developmental processes and immunity pathways are expanded in taller plants. As taller plants also have longer generation times, we hypothesize that the expansion of components of the immune system in these species may a consequence of the selective pressure caused by chronic pathogenic infections, a problem likely to be far more critical for long-living species that may take years before achieving fertility.

**Estimation of ancestor state for maximum height in Angiosperms**

We estimated the ancestral height state for all the internal nodes of the phylogeny based on the height values of the 54 extant species with high-quality genomes available (see Material and Methods, section “Estimation of ancestor states for heigh in Angiosperms”). Due to the high degree of height variation observed, most internal nodes were given average values, and, based on visual inspection, we found no evolutionary trend on height variation across the distinct angiosperm lineages (Supplementary Figure 5B). On the contrary, we notice the increase and decrease in plant height to be spread across different clades. These results indicate that many independent events of increase and decrease in plant height have occurred in the evolutionary history of flowering plants, suggesting that distinct evolutionary strategies under diverse selective pressures might explain the height variation in of angiosperms (e.g., higher longevity in taller species vs annual plants in shorter species) (Lanfear et al., 2013).

**CALANGO package and dependencies**

The CALANGO package is designed as an open-source tool for comparative genomics. CALANGO was developed as a CRAN-compliant R package (R_Development_Core_Team, 2016) and makes full use of the existing R ecosystem to implement its internal routines using code that has been validated by other researchers and developers. Our tool is implemented as an open-source R package distributed through the Comprehensive R Archive Network (CRAN). The package outputs results in the form of fully-functional dynamical HTML5 sites containing several summary statistics and other useful quantities, as well as interactive visual representations. CALANGO also outputs all results and reproducibility parameters as a list object, allowing easy integration with other bioinformatics pipelines (Fig. 1C).

CALANGO uses several R libraries to handle the different data types needed. The CALANGO analysis routines import functions from packages *ape* (Paradis and Schliep, 2019) (to read nexus and newick phylogenetic trees and resolve multichotomies, and to calculate phylogeny-independent contrasts and the correlation structures arising from phylogenetic relationships); *taxize* (Chamberlain and Szocs, 2013) (to retrieve and process taxonomical hierarchies); *GO.db* (Carlson, 2016) and *AnnotationDbi* (Hervé Pagès, 2020) (to process GO annotation data); *KEGGREST* (Tenenbaum, 2020) (to process KEGG databases); and *nlme* (Pinheiro J, 2021) (to fit models using generalized least squares). CALANGO also imports functions from several packages to compose its visual output, namely: *dendextend* (Galili, 2015), *rmarkdown* (JJ Allaire and Ushey, 2020) *heatmaply* (Galili et al., 2018), *ggplot2* (Wickham, 2020), *plotly* (Sievert, 2020), *DT* (Yihui Xie, 2021), *htmltools* (Joe Cheng, 2021) and *htmlwidgets* (Ramnath Vaidyanathan, 2020). Other general-purpose packages used within CALANGO are *pbmcapply* (Kevin Kuang, 2019) (for progress bars when using parallel processing); *assertthat* (Wickham, 2019) (for input verification); *BiocManager* (Morgan, 2019) (to retrieve and update dependencies from Bioconductor, namely *KEGGREST*, *GO.db* and *AnnotateDbi*); and *pkgdown* (Hadley Wickham, 2019) (to automatically generate the project home page). Package updates on the CALANGO repository are automatically verified using Github Actions on the latest R versions for Windows and Mac OS, as well as for both the release and devel R versions on Ubuntu 20.04 LTS, to ensure code integrity. Future versions of CALANGO are planned to reduce the number of distinct dependencies so as to make the tool more resilient to changes in external package functionalities.

**SUPPLEMENTARY REFERENCES**

Arndt, D., Grant, J.R., Marcu, A., Sajed, T., Pon, A., Liang, Y., and Wishart, D.S. (2016). PHASTER: a better, faster version of the PHAST phage search tool. Nucleic acids research *44*, W16-21.

Barr, J.J., Auro, R., Furlan, M., Whiteson, K.L., Erb, M.L., Pogliano, J., Stotland, A., Wolkowicz, R., Cutting, A.S., Doran, K.S.*, et al.* (2013). Bacteriophage adhering to mucus provide a non-host-derived immunity. Proceedings of the National Academy of Sciences of the United States of America *110*, 10771-10776.

Bertozzi Silva, J., Storms, Z., and Sauvageau, D. (2016). Host receptors for bacteriophage adsorption. FEMS Microbiol Lett *363*.

Carlson, M. (2016). GO.db: A set of annotation maps describing the entire Gene Ontology.

Chamberlain, S.A., and Szocs, E. (2013). taxize: taxonomic search and retrieval in R. F1000Research *2*, 191.

Cirillo, D.M., Heffernan, E.J., Wu, L., Harwood, J., Fierer, J., and Guiney, D.G. (1996). Identification of a domain in Rck, a product of the Salmonella typhimurium virulence plasmid, required for both serum resistance and cell invasion. Infect Immun *64*, 2019-2023.

Cotter, P.D., Ross, R.P., and Hill, C. (2013). Bacteriocins - a viable alternative to antibiotics? Nat Rev Microbiol *11*, 95-105.

Dedrick, R.M., Jacobs-Sera, D., Bustamante, C.A., Garlena, R.A., Mavrich, T.N., Pope, W.H., Reyes, J.C., Russell, D.A., Adair, T., Alvey, R.*, et al.* (2017). Prophage-mediated defence against viral attack and viral counter-defence. Nat Microbiol *2*, 16251.

Dufayard, J.F., Bettembourg, M., Fischer, I., Droc, G., Guiderdoni, E., Perin, C., Chantret, N., and Dievart, A. (2017). New Insights on Leucine-Rich Repeats Receptor-Like Kinase Orthologous Relationships in Angiosperms. Frontiers in plant science *8*, 381.

Ehrbar, K., and Hardt, W.D. (2005). Bacteriophage-encoded type III effectors in Salmonella enterica subspecies 1 serovar Typhimurium. Infect Genet Evol *5*, 1-9.

Fernandez, L., Rodriguez, A., and Garcia, P. (2018). Phage or foe: an insight into the impact of viral predation on microbial communities. The ISME journal *12*, 1171-1179.

Galili, T. (2015). dendextend: an R package for visualizing, adjusting and comparing trees of hierarchical clustering. Bioinformatics *31*, 3718-3720.

Galili, T., O'Callaghan, A., Sidi, J., and Sievert, C. (2018). heatmaply: an R package for creating interactive cluster heatmaps for online publishing. Bioinformatics *34*, 1600-1602.

Gordillo Altamirano, F., Forsyth, J.H., Patwa, R., Kostoulias, X., Trim, M., Subedi, D., Archer, S.K., Morris, F.C., Oliveira, C., Kielty, L.*, et al.* (2021). Bacteriophage-resistant Acinetobacter baumannii are resensitized to antimicrobials. Nat Microbiol *6*, 157-161.

Hadley Wickham, J.H. (2019). pkgdown: Make Static HTML Documentation for a Package.

Hervé Pagès, M.C., Seth Falcon, Nianhua Li (2020). AnnotationDbi: Manipulation of SQLite-based annotations in Bioconductor.

Jarvis, K.G., Giron, J.A., Jerse, A.E., McDaniel, T.K., Donnenberg, M.S., and Kaper, J.B. (1995). Enteropathogenic Escherichia coli contains a putative type III secretion system necessary for the export of proteins involved in attaching and effacing lesion formation. Proceedings of the National Academy of Sciences of the United States of America *92*, 7996-8000.

JJ Allaire, Y.X., Jonathan McPherson, Javier Luraschi, Kevin, and Ushey, A.A., Hadley Wickham, Joe Cheng, Winston Chang, Richard Iannone (2020). rmarkdown: Dynamic Documents for R.

Joe Cheng, C.S., Winston Chang, Yihui Xie, Jeff Allen (2021). htmltools: Tools for HTML.

Jones, P., Binns, D., Chang, H.Y., Fraser, M., Li, W., McAnulla, C., McWilliam, H., Maslen, J., Mitchell, A., Nuka, G.*, et al.* (2014). InterProScan 5: genome-scale protein function classification. Bioinformatics *30*, 1236-1240.

Kevin Kuang, Q.K., Francesco Napolitano (2019). pbmcapply: Tracking the Progress of Mc*pply with Progress Bar.

.

Kumar, V., Donev, E.N., Barbut, F.R., Kushwah, S., Mannapperuma, C., Urbancsok, J., and Mellerowicz, E.J. (2020). Genome-Wide Identification of Populus Malectin/Malectin-Like Domain-Containing Proteins and Expression Analyses Reveal Novel Candidates for Signaling and Regulation of Wood Development. Frontiers in plant science *11*, 588846.

Kumar, V., Hainaut, M., Delhomme, N., Mannapperuma, C., Immerzeel, P., Street, N.R., Henrissat, B., and Mellerowicz, E.J. (2019). Poplar carbohydrate-active enzymes: whole-genome annotation and functional analyses based on RNA expression data. Plant J *99*, 589-609.

Lanfear, R., Ho, S.Y., Jonathan Davies, T., Moles, A.T., Aarssen, L., Swenson, N.G., Warman, L., Zanne, A.E., and Allen, A.P. (2013). Taller plants have lower rates of molecular evolution. Nat Commun *4*, 1879.

Leon, M., and Bastias, R. (2015). Virulence reduction in bacteriophage resistant bacteria. Frontiers in microbiology *6*, 343.

Maslov, S., and Sneppen, K. (2017). Population cycles and species diversity in dynamic Kill-the-Winner model of microbial ecosystems. Scientific reports *7*, 39642.

Morgan, M. (2019). BiocManager: Access the Bioconductor Project Package Repository.

Mosavi, L.K., Cammett, T.J., Desrosiers, D.C., and Peng, Z.Y. (2004). The ankyrin repeat as molecular architecture for protein recognition. Protein Sci *13*, 1435-1448.

Nishiyama, T., Noguchi, H., Yoshida, H., Park, S.Y., and Tame, J.R. (2013). The structure of the deacetylase domain of Escherichia coli PgaB, an enzyme required for biofilm formation: a circularly permuted member of the carbohydrate esterase 4 family. Acta Crystallogr D Biol Crystallogr *69*, 44-51.

Paradis, E., and Schliep, K. (2019). ape 5.0: an environment for modern phylogenetics and evolutionary analyses in R. Bioinformatics *35*, 526-528.

Park, B.S., and Lee, J.O. (2013). Recognition of lipopolysaccharide pattern by TLR4 complexes. Exp Mol Med *45*, e66.

Pawluk, A., Bondy-Denomy, J., Cheung, V.H., Maxwell, K.L., and Davidson, A.R. (2014). A new group of phage anti-CRISPR genes inhibits the type I-E CRISPR-Cas system of Pseudomonas aeruginosa. mBio *5*, e00896.

Pinard, D., Mizrachi, E., Hefer, C.A., Kersting, A.R., Joubert, F., Douglas, C.J., Mansfield, S.D., and Myburg, A.A. (2015). Comparative analysis of plant carbohydrate active enZymes and their role in xylogenesis. BMC genomics *16*, 402.

Pinheiro J, B.D., DebRoy S, Sarkar D, R Core Team (2021). nlme: Linear and Nonlinear Mixed Effects Models.

Pulkkinen, W.S., and Miller, S.I. (1991). A Salmonella typhimurium virulence protein is similar to a Yersinia enterocolitica invasion protein and a bacteriophage lambda outer membrane protein. J Bacteriol *173*, 86-93.

R_Development_Core_Team (2016). R: A Language and Environment for Statistical Computing.

Ramisetty, B.C.M., and Sudhakari, P.A. (2019). Bacterial 'Grounded' Prophages: Hotspots for Genetic Renovation and Innovation. Front Genet *10*, 65.

Ramnath Vaidyanathan, Y.X., JJ Allaire, Joe Cheng, Carson Sievert, Kenton Russell (2020). htmlwidgets: HTML Widgets for R.

Salzberg, S.L. (2019). Next-generation genome annotation: we still struggle to get it right. Genome biology *20*, 92.

Schallus, T., Jaeckh, C., Feher, K., Palma, A.S., Liu, Y., Simpson, J.C., Mackeen, M., Stier, G., Gibson, T.J., Feizi, T.*, et al.* (2008). Malectin: a novel carbohydrate-binding protein of the endoplasmic reticulum and a candidate player in the early steps of protein N-glycosylation. Mol Biol Cell *19*, 3404-3414.

Sievert, C. (2020). Interactive Web-Based Data Visualization with R, plotly, and shiny, 1 edn (Chapman and Hall/CRC).

Silveira, C.B., and Rohwer, F.L. (2016). Piggyback-the-Winner in host-associated microbial communities. NPJ Biofilms Microbiomes *2*, 16010.

Simmons, E.L., Bond, M.C., Koskella, B., Drescher, K., Bucci, V., and Nadell, C.D. (2020). Biofilm Structure Promotes Coexistence of Phage-Resistant and Phage-Susceptible Bacteria. mSystems *5*.

Steyert, S.R., and Kaper, J.B. (2012). Contribution of urease to colonization by Shiga toxin-producing Escherichia coli. Infect Immun *80*, 2589-2600.

Tenenbaum, D. (2020). KEGGREST: Client-side REST access to KEGG.

Tuskan, G.A., Difazio, S., Jansson, S., Bohlmann, J., Grigoriev, I., Hellsten, U., Putnam, N., Ralph, S., Rombauts, S., Salamov, A.*, et al.* (2006). The genome of black cottonwood, Populus trichocarpa (Torr. & Gray). Science (New York, NY *313*, 1596-1604.

Vandecraen, J., Chandler, M., Aertsen, A., and Van Houdt, R. (2017). The impact of insertion sequences on bacterial genome plasticity and adaptability. Crit Rev Microbiol *43*, 709-730.

Wickham, H. (2019). assertthat: Easy Pre and Post Assertions.

Wickham, H. (2020). ggplot2: Elegant Graphics for Data Analysis, 3 edn (Springer International Publishing).

Wu, B., Skarina, T., Yee, A., Jobin, M.C., Dileo, R., Semesi, A., Fares, C., Lemak, A., Coombes, B.K., Arrowsmith, C.H.*, et al.* (2010). NleG Type 3 effectors from enterohaemorrhagic Escherichia coli are U-Box E3 ubiquitin ligases. PLoS pathogens *6*, e1000960.

Xiao, Y., Stegmann, M., Han, Z., DeFalco, T.A., Parys, K., Xu, L., Belkhadir, Y., Zipfel, C., and Chai, J. (2019). Mechanisms of RALF peptide perception by a heterotypic receptor complex. Nature *572*, 270-274.

Yang, H., Wang, D., Guo, L., Pan, H., Yvon, R., Garman, S., Wu, H.M., and Cheung, A.Y. (2021). Malectin/Malectin-like domain-containing proteins: A repertoire of cell surface molecules with broad functional potential. Cell Surf *7*, 100056.

Yihui Xie, J.C., Xianying Tan (2021). DT: A Wrapper of the JavaScript Library 'DataTables'.

**SUPPLEMENTARY FIGURE AND TABLE LEGENDS**

**Supplementary Table 1. *Escherichia coli* genomic and phenotypic data.**

**Supplementary Table 2. Protein domains and Gene Ontology terms associated with prophage density in the *E. coli* dataset.**

**Supplementary Table 3. Angiosperm genomic and phenotypic data.**

**Supplementary Table 4. *Arabidopsis thaliana* genes and protein domains positively associated with maximum height in Angiosperms.**

**Supplementary Figure 1. Genomic components, annotation terms, and feature normalization.** A) Genomic components and their annotation terms: 1) Protein-centered annotation to Gene Ontology terms. A single protein-coding gene may be annotated to as many GO terms as needed. 2) Domain-centered annotation of protein domains to domain IDs, such as the ones available in Pfam. 3) Promoter-centered annotation of distinct binding sites of specific DNA binding proteins. 4) relationship between annotation term IDs and their definitions (in this case, hypothetical protein domain IDs and their biological roles). B) Hypothetical genomes and their associated metadata for normalization. The figure represents annotation terms with either the same number of copies in all genomes (orange boxes) or with a variable number of copies in all genomes (blue boxes). The table contains QVAL for these genomes, the normalization factor for annotation counts (sum of all annotation terms), raw annotation term counts, and their relative frequencies (raw counts of each annotation term in a genome normalized by the sum of all annotation terms in a genome). C) Correlation of QVAL, count, and frequency data. Each plot contains the correlation for the hypothetical variables in Supplementary Figure 1B. It is possible to observe that, the annotation term represented by blue boxes is associated with QVAL (both count and normalized values). The orange term, however, is not associated when considering count data, as it is present as a single-copy region in all genomes. As the relative frequencies of the orange annotation terms in each genome are dependent on the relative frequency of blue annotation terms, a relative increase in the frequency of blue annotation terms causes the relative decrease of orange annotation terms in the same genome.

**Supplementary Figure 2 - Heatmap of protein domains associated with prophage density in *E. coli* after the removal of genes of viral origin as produced by CALANGO.** Species clustering was performed using the *E. coli* phylogeny produced in this study, annotation terms clustering was performed using Manhattan distance and average clustering. Red arrows indicate *E. coli* lineages with some of the highest values of domains of viral origin (before removal of genes of viral origin) and virulence factors.

**Supplementary Figure 3 – Additional examples of Pfam domains associated with prophage density.** For each Pfam domain ID, from left to right, CALANGO provides two plots for traditional association statistics (a scatterplot, together with a linear model, for direct data visualization, and a scatterplot of ranked data, together with a locally estimated scatterplot smoothing (LOESS)) and a plot containing the contrast data for phylogeny-aware linear regression of phylogenetically independent contrasts.

**Supplementary Figure 4 – Additional examples of GO terms associated with prophage density.** For each GO term ID, from left to right, CALANGO provides two plots for traditional association statistics (a scatterplot, together with a linear model, for direct data visualization, and a scatterplot of ranked data, together with a locally estimated scatterplot smoothing (LOESS)) and a plot containing the contrast data for phylogeny-aware linear regression of phylogenetically independent contrasts.

**Supplementary Figure 5 – Evolution of maximum height in Angiosperms.** A) Phenotypic variation of maximum height in the 54 Angiosperm species used in this work. B) Ancestral state reconstruction for maximum height.

**Supplementary Figure 6 – Additional examples of Pfam terms associated with maximum height in Angiosperms.** From left to right: linear models from raw count data, phylogeny-aware linear models; boxplot with raw counts; boxplot with normalized counts.
