## Supplementary figures and images for "CALANGO: a phylogeny-aware comparative genomics tool for discovering quantitative genotype-phenotype associations across species"

### Supplementary Figure 1

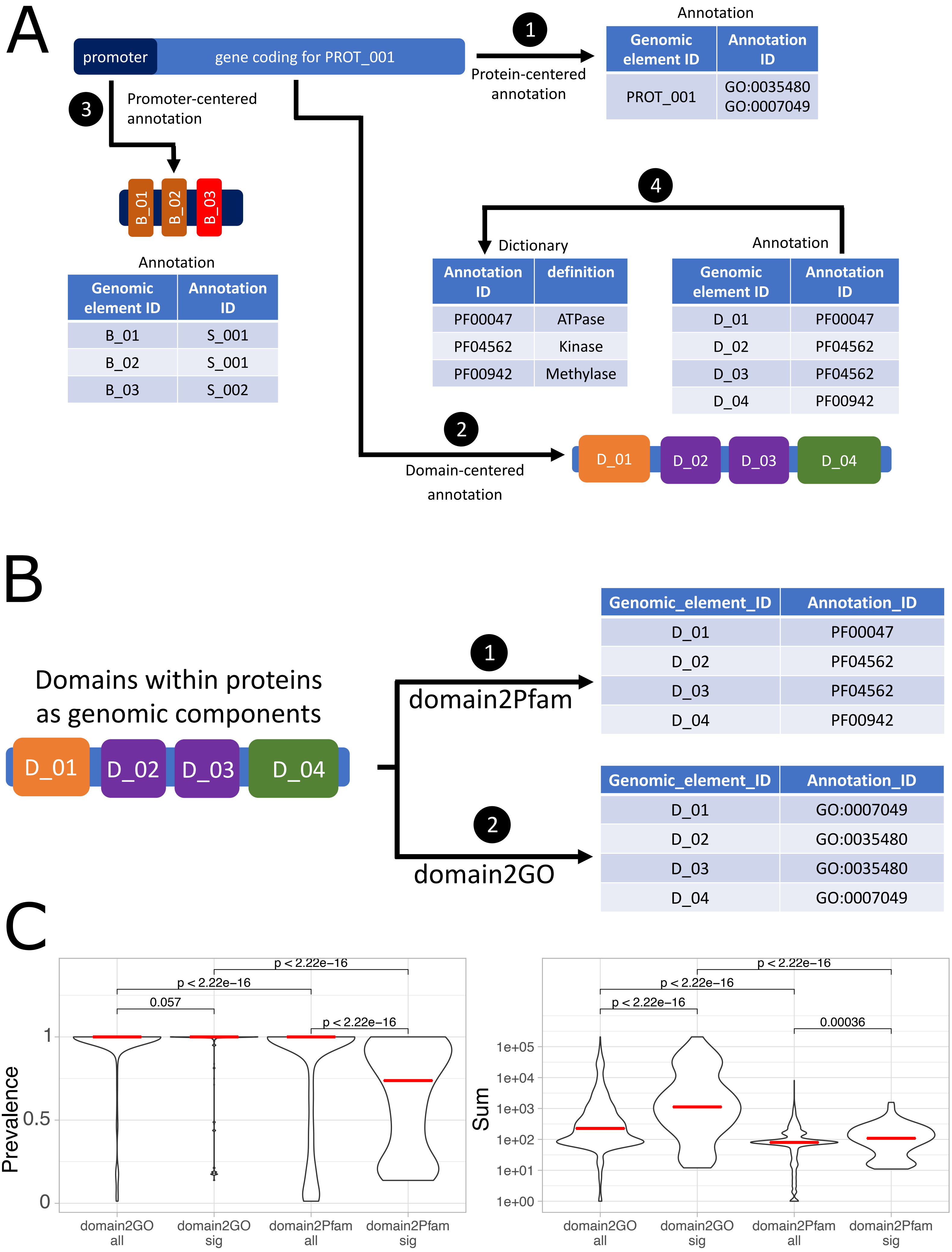

### Supplementary Figure 3

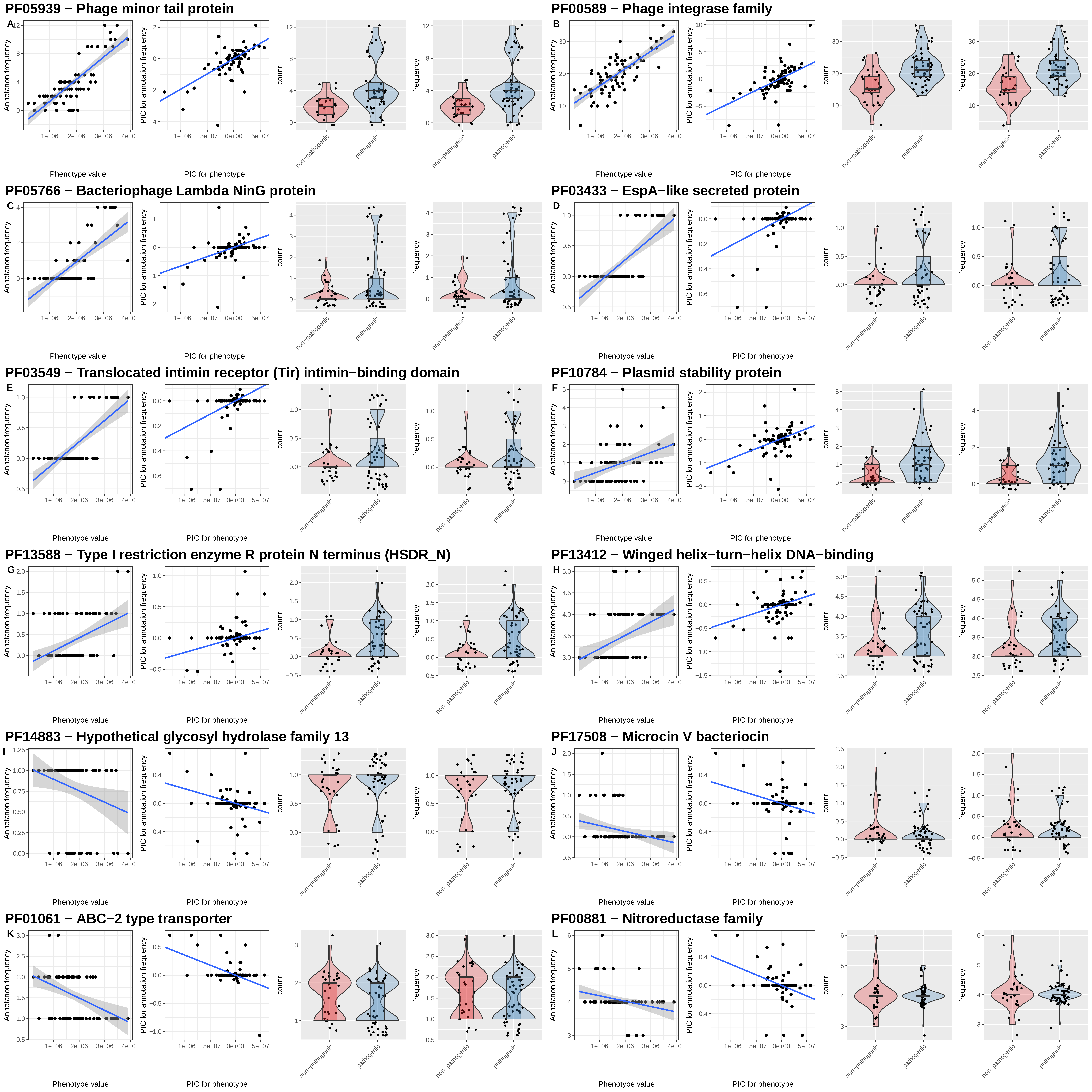

### Supplementary Figure 4

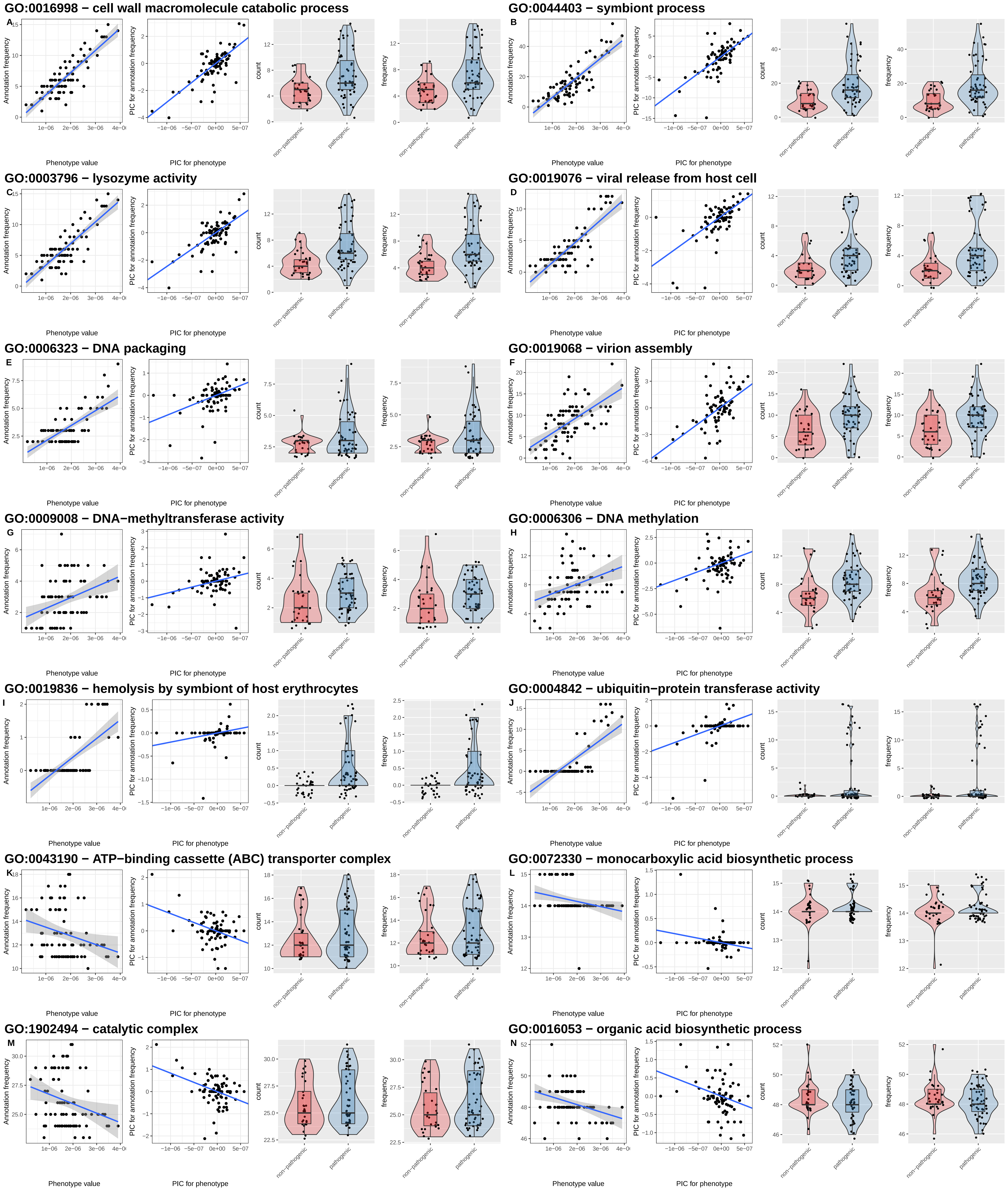

### Supplementary Figure 5

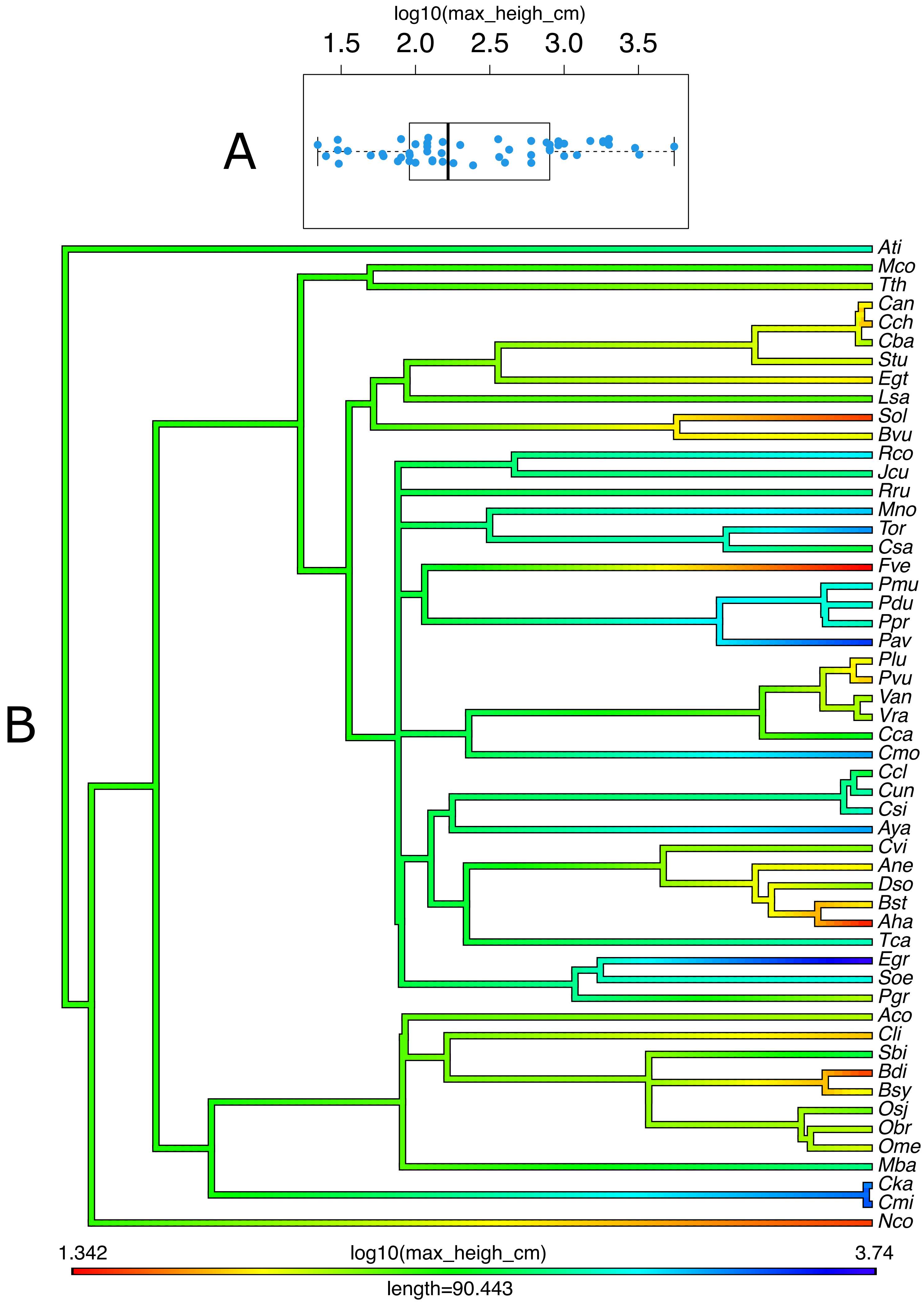

### Supplementary Figure 6

#### PF00560 − Leucine Rich Repeat

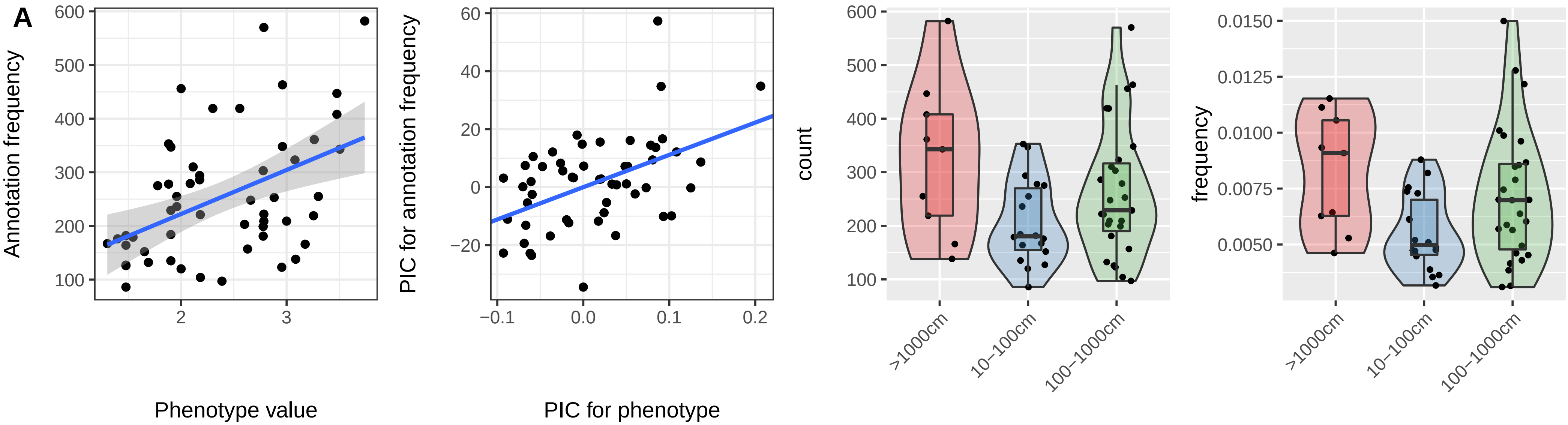


#### PF08276 − PAN−like domain

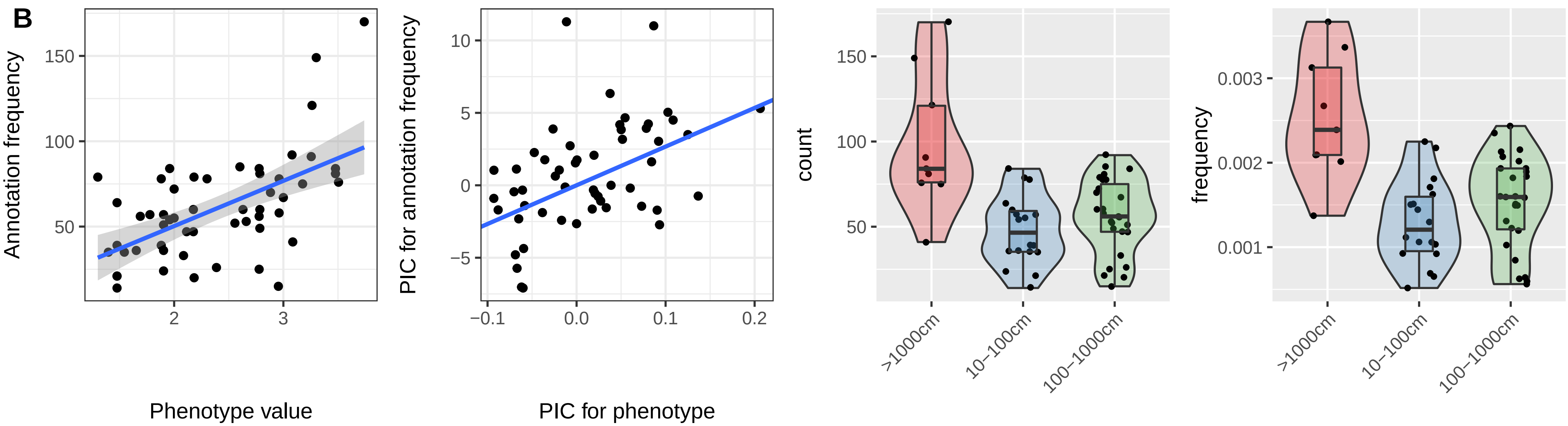


#### PF11883 − Domain of unknown function (DUF3403)

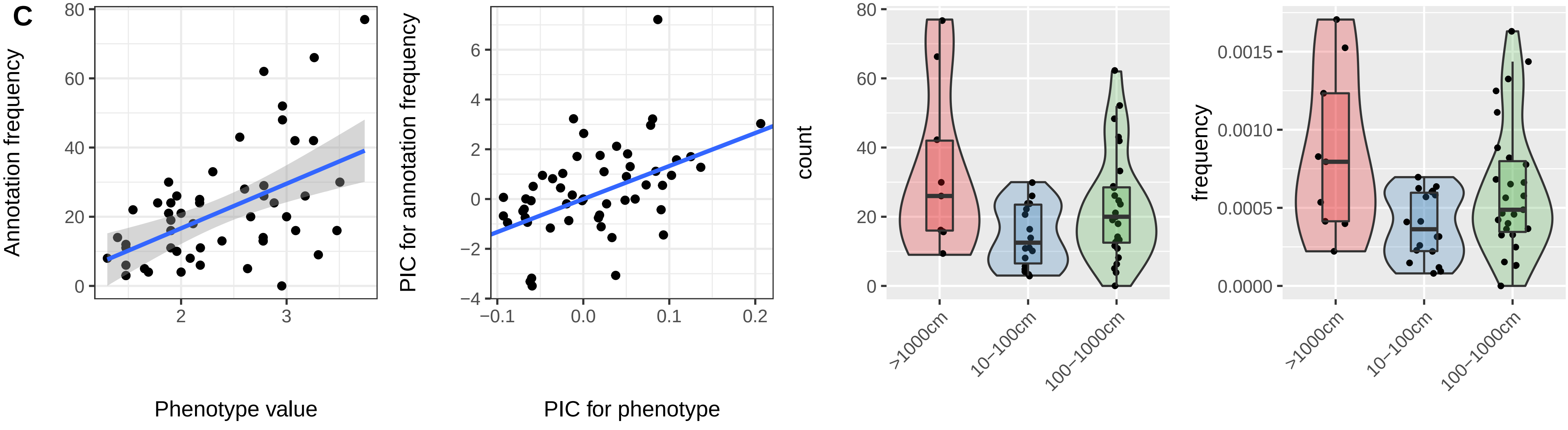
