## Supplementary Figure 2 for "CALANGO: a phylogeny-aware comparative genomics tool for discovering quantitative genotype-phenotype associations across species"

A

**Genomes**

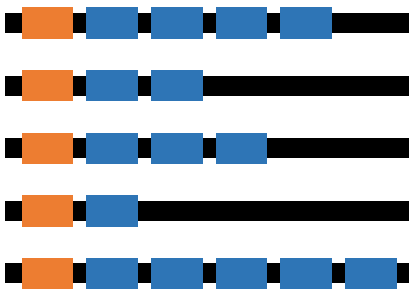

| Total annotation terms | QVAL | Blue count | Orange count | Blue Frequency | Orange frequency |
|---|---|---|---|---|---|
| 5 | 0.1 | 4 | 1 | 0.8 | 0.2 |
| 3 | 0.25 | 2 | 1 | 0.67 | 0.33 |
| 4 | 0.15 | 3 | 1 | 0.75 | 0.25 |
| 2 | 0.4 | 1 | 1 | 0.5 | 0.5 |
| 6 | 0 | 5 | 1 | 0.83 | 0.17 |

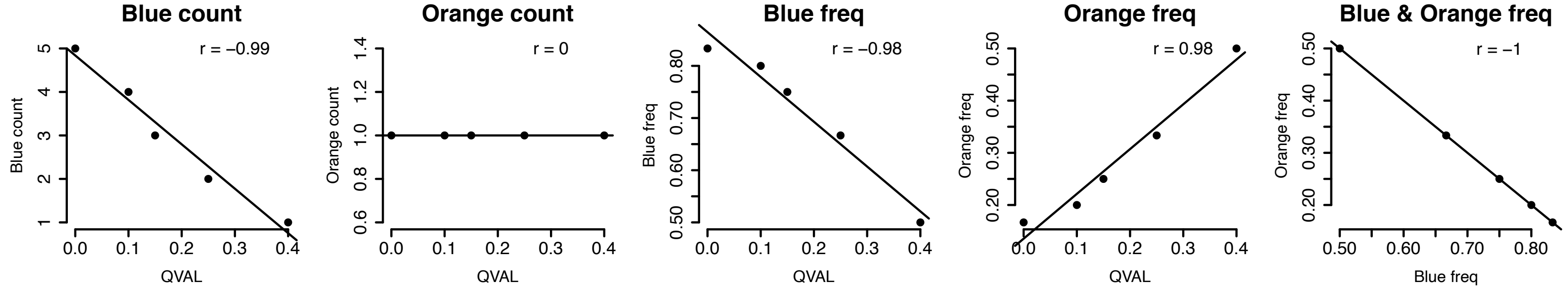


B

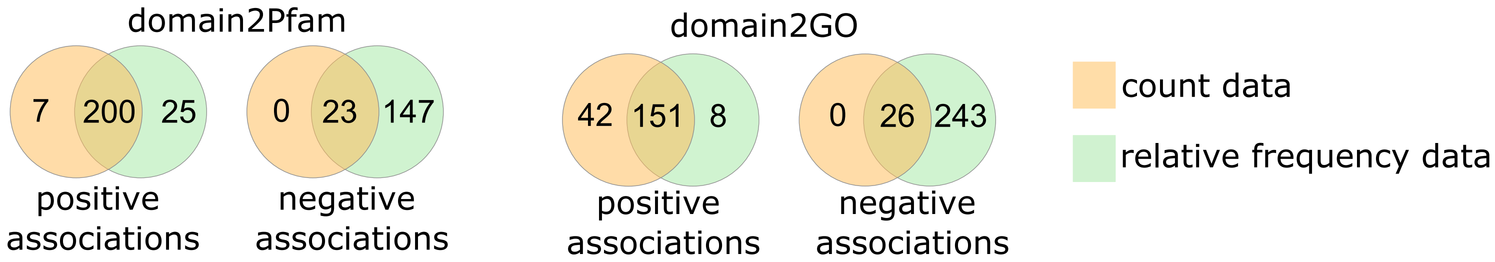
